# Perturbation of *de novo* lipogenesis hinders MERS-CoV assembly and release, but not the biogenesis of viral replication organelles

**DOI:** 10.1101/2024.08.21.608937

**Authors:** M. Soultsioti, A.W.M. de Jong, N. Blomberg, A. Tas, M. Giera, E. J. Snijder, M. Barcena

## Abstract

Coronaviruses hijack host cell metabolic pathways and resources to support their replication. They induce extensive host endomembrane remodeling to generate viral replication organelles and exploit host membranes for assembly and budding of their enveloped progeny virions. Because of the overall significance of host membranes, we sought to gain insight into the role of host factors involved in lipid metabolism in cells infected with Middle East respiratory syndrome coronavirus (MERS-CoV). We employed a single-cycle infection approach in combination with pharmacological inhibitors, biochemical assays, lipidomics, light and electron microscopy. Pharmacological inhibition of acetyl-CoA carboxylase (ACC) and fatty acid synthase (FASN), key host factors in *de novo* fatty acid biosynthesis, led to pronounced inhibition of MERS-CoV particle release. Inhibition of ACC led to a profound metabolic switch in Huh7 cells, altering their lipidomic profile and inducing lipolysis. However, despite the extensive changes induced by the ACC inhibitor, the biogenesis of viral replication organelles remained unaffected. Instead, ACC inhibition appeared to affect the trafficking and post-translational modifications of the MERS-CoV envelope proteins. Electron microscopy revealed an accumulation of nucleocapsids in early budding stages, indicating that MERS-CoV assembly is adversely impacted by ACC inhibition. Notably, inhibition of palmitoylation resulted in similar effects, while supplementation of exogenous palmitic acid reversed the compound’s inhibitory effects, possibly reflecting a crucial need for palmitoylation of the MERS-CoV Spike and Envelope proteins for their role in virus particle assembly.

**Importance:** Middle East respiratory syndrome coronavirus (MERS-CoV) is the etiological agent of a zoonotic respiratory disease of limited transmissibility between humans. However, MERS-CoV is still considered a high-priority pathogen and is closely monitored by WHO due to its high lethality rate of around 35% of laboratory-confirmed infections. Like other positive-strand RNA viruses, MERS-CoV relies on the host cell’s endomembranes to support various stages of its replication cycle. However, in spite of this general reliance of MERS-CoV replication on host cell lipid metabolism, mechanistic insights are still very limited. In our study, we show that pharmacological inhibition of acetyl-CoA carboxylase (ACC), a key enzyme in the host cell’s fatty acid biosynthesis pathway, significantly disrupts MERS-CoV particle assembly without exerting a negative effect on the biogenesis of viral replication organelles. Furthermore, our study highlights the potential of ACC as a target for the development of host-directed antiviral therapeutics against coronaviruses.

## Introduction

Viruses rely heavily on the host cell’s metabolic and biosynthetic capabilities to support their replication cycle. As such, the manipulation of host cell metabolic pathways has been a long-standing research topic for a range of evolutionary divergent DNA and RNA virus families, such as herpesviruses, flaviviruses, and picornaviruses, as reviewed in (1, 2). In the case of coronaviruses, virus-host interactions related to cellular metabolism constituted a rather poorly studied area until the emergence of severe acute respiratory syndrome coronavirus-2 (SARS-CoV-2) in December 2019 (3, 4). Since then, several studies have highlighted that coronaviruses, like other virus families, hijack host metabolic pathways to support various stages of their replication (5–12).

During their replication cycle (reviewed in (13–15)), coronaviruses, including the highly pathogenic SARS-CoV, SARS-CoV-2, and Middle East respiratory syndrome coronavirus (MERS-CoV), hijack intracellular membranes to accommodate viral RNA synthesis (16–23) and the assembly of enveloped progeny virions (21, 24–27). Briefly, after coronavirus entry and release of the positive-strand RNA genome (gRNA) into the cytoplasm, the gRNA is translated into two large replicase polyproteins. These are autoproteolytically processed by internally encoded viral proteases, resulting in the generation of 16 viral nonstructural proteins (nsps) that engage in viral RNA synthesis and a range of virus-host interactions. All coronaviruses studied so far induce the formation of a perinuclear network of modified membrane structures that derive from the endoplasmic reticulum (ER). This network includes double-membrane vesicles (DMVs), paired or convoluted membranes (CMs), and double-membrane spherules (DMSs), which are collectively referred to as the coronavirus replication organelle (RO) (18, 20). DMVs are the most abundant structures and constitute the primary site of viral genome replication and generation of subgenomic (sg) mRNAs (20, 21). A molecular pore complex was found to span the DMV’s double membrane, thus connecting the DMV interior with the cytosol (28–30) and presumably constituting a gateway for viral RNA export. In the cytoplasm, the sg mRNAs are translated, giving rise to the viral structural and accessory proteins. In parallel, part of the newly synthesized gRNAs associate with Nucleocapsid (N) proteins to form ribonucleoprotein (RNP) complexes, which constitute the nucleocapsid substructures that will be packaged in the virus particle. During their synthesis, the viral envelope proteins associate with membranes of the secretory pathway, into which the nucleocapsid buds. Progeny virions acquire their envelope from intracellular membranes that primarily originate from the ER-Golgi intermediate compartment (ERGIC) (21, 26), after which they are released from the infected cell. Historically, viral egress was thought to occur via the biosynthetic secretory pathway (31), but recent evidence suggests that coronaviruses may (also) employ parts of the lysosomal pathway (27, 32–34).

Given the overall significance of host cell lipids and membranes in pivotal stages of coronavirus replication, it is not surprising that recent genome-wide association studies (GWAS) have identified host factors involved in fatty acid biosynthesis/*de novo* lipogenesis (DNL) and cholesterol biosynthesis to be important for the replication of SARS-CoV-2 and other coronaviruses (35–37). Host factors identified included the sterol regulatory element-binding proteins (SREBP-1 and SREBP-2) and SREBP cleavage-activating protein (SCAP), which are major regulators of DNL and cholesterol biosynthesis (38). A prior study also implicated SREBPs in MERS-CoV replication, potentially in DMV formation (6). Due to the impact of the SARS-CoV-2 pandemic and the urgency of developing broad-spectrum antiviral therapeutics, the potential of enzymes of the DNL pathway as drug targets for intervention and treatment of COVID-19 has been explored, yielding some promising results (10, 39, 40). It should however be noted that, while most studies on coronavirus replication so far have uncovered a general reliance on DNL-related pathways, further mechanistic insights remain scarce.

The aim of our present study was to explore the role of DNL in the MERS-CoV replication cycle. Our results show that inhibition of acetyl-CoA carboxylase (ACC) disrupts the assembly of infectious MERS-CoV progeny, while the formation of viral replication organelles remains unaffected. The inhibitory effect on virus assembly could be partially reversed by supplementation with exogenous palmitic acid, possibly reflecting a need for palmitoylation of the MERS-CoV Spike (S) and Envelope (E) proteins. Our results also highlight the potential of ACC as a target for the development of host-directed antiviral treatment.

## Results

### MERS-CoV infection increases gene expression of host factors involved in DNL

*De novo* lipogenesis begins with the generation of acetyl-CoA from citrate through the activity of ATP citrate lyase (ACLY) (Figure 1A). Acetyl-CoA is further carboxylated by the rate-limiting enzyme ACC, leading to the production of malonyl-CoA. Malonyl-CoA then serves as a substrate for the fatty acid synthase enzyme (FASN), leading to the generation of saturated fatty acids (such as myristic acid, palmitic acid, and stearic acid) that can be utilized in post-translational modifications (e.g., myristoylation and palmitoylation), phospholipid synthesis, and fatty acid catabolism (β-oxidation) (Figure 1A).

**Figure 1.**
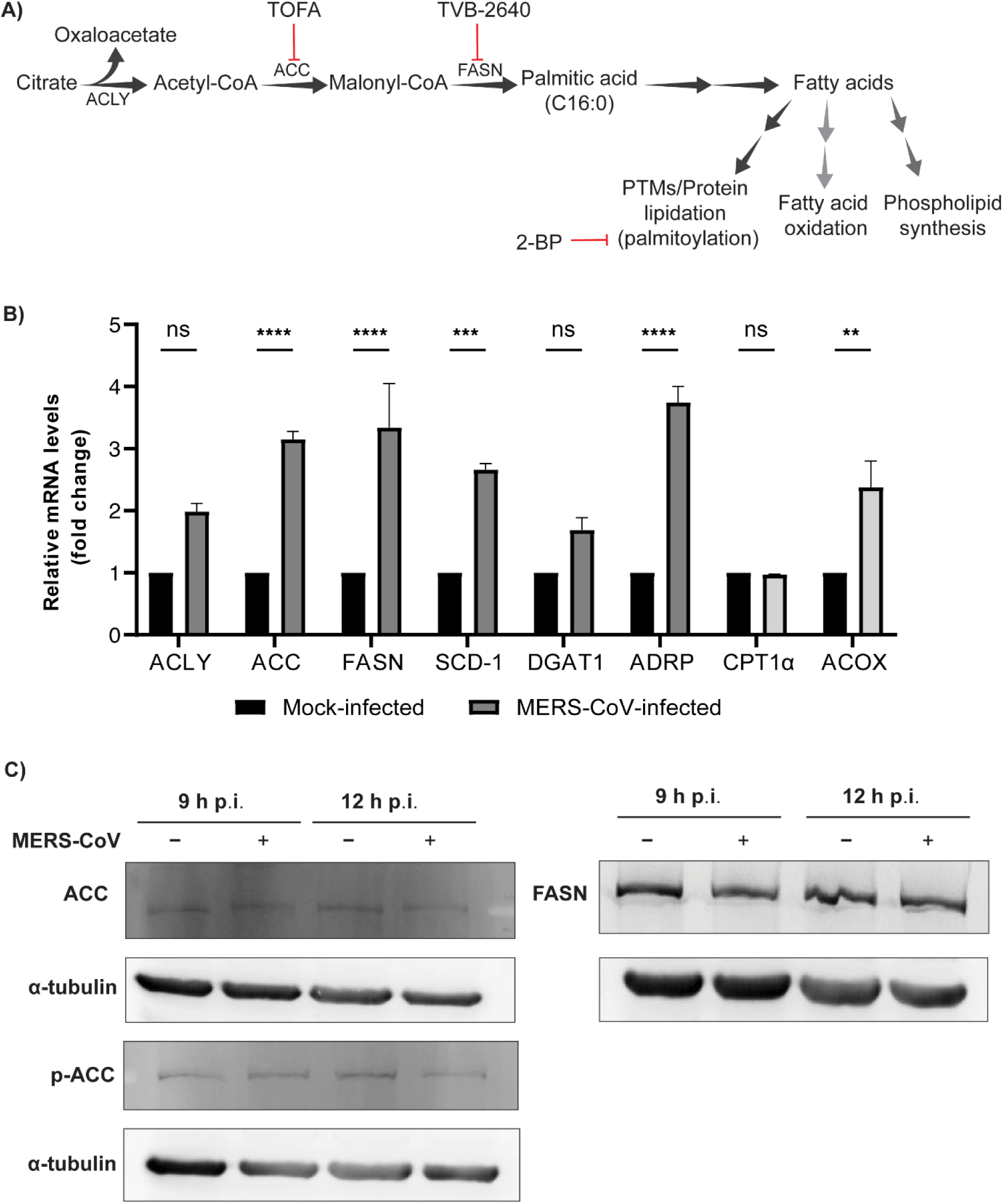
Gene expression of host factors of the DNL pathway increases upon MERS-CoV infection. A) Schematic representation of the main steps of the *de novo* lipogenesis pathway, the key enzymes involved (ACLY, ACC, FASN), and the downstream biological functions of the resulting fatty acids. Double arrows indicate that multiple reactions are involved but not depicted here. Small-molecule inhibitors used in this study are denoted in red. PTMs: posttranslational modifications. B) Gene expression analysis of key host factors participating in DNL and phospholipid synthesis (dark gray) or fatty acid oxidation (light gray) in mock-infected Huh7 cells in comparison with MERS-CoV-infected cells (MOI 5) at 12 h p.i. The mRNA levels of *ACLY, ACC, FASN, SCD-1, DGAT1, ADRP, CPT1a* and *ACOX* were determined by RT-qPCR and normalized against the mock-infected control. C) Western Blot analysis of ACC, phospho-ACC and FASN in MERS-CoV infected Huh7 cells, at 9 and 12 h p.i. Data are represented as mean ± SD of two biological replicates. Statistical significance was calculated using two-way ANOVA and applying Bonferroni multiple comparison correction, * p<0.05, ** p< 0.01, *** p< 0.001, **** p< 0.0001.

First, we investigated if MERS-CoV infection altered the gene expression levels of various host factors involved in DNL (*ACLY, ACC, FASN, SCD-1*), phospholipid synthesis (*DGAT1*), formation of lipid droplets (LDs) (*ADRP*), or fatty acid oxidation pathways (*CPT1α* and *ACOX*). To this end, Huh7 cells were infected with MERS-CoV at a high multiplicity of infection (MOI 5). At 12 h p.i., cells were lysed and mRNA expression was analyzed by RT-qPCR. This timepoint was chosen as it combined abundant viral RNA synthesis with the early phase of virus assembly and release. Under such conditions, we observed an increase in the mRNA levels of host factors involved in initial steps of the DNL pathway, such as the rate-limiting enzymes ACC and FASN, and an increase in the mRNA levels of ADRP (Figure 1B). From the fatty acid oxidation pathway factors that were analyzed, only ACOX was moderately increased. We then evaluated the expression of ACC and FASN on a protein level by Western blot analysis (Figure 1C). Despite the observed alterations in mRNA levels, a clear change in the protein abundance of ACC and FASN could not be detected. However, this result may be attributed to the limited sensitivity of the assay or a reduced turn-over of these proteins. During high energy requirement, the activity of ACC is regulated via its phosphorylation by AMP-activated protein kinase (AMPK), which leads to inactivation of ACC. In this context, we also evaluated the levels of phospho-ACC by Western blot, but did not observe any changes (Figure 1C), indicating that ACC’s activity is not inhibited in MERS-CoV-infected cells. Our combined observations suggested that host factors of the DNL branch of lipid metabolism are transcriptionally upregulated in MERS-CoV-infected cells.

### DNL inhibition does not affect viral RNA synthesis but impairs a later step in the MERS-CoV replication cycle

To investigate the role of DNL pathways in coronavirus replication, we made use of the fast-acting and potent small-molecule inhibitor 5-(tetradecyloxy)-2-furoic acid (TOFA), which blocks the activity of ACC and thus the synthesis of fatty acids (Figure 1A) (41, 42). Our aim was to explore whether pharmacological inhibition of ACC would influence viral replication, in particular the steps that directly involve host cell endomembranes: replication organelle formation and assembly of progeny virions. For this, Huh7 cells were infected with MERS-CoV at MOI 5 and treated with non-cytotoxic concentrations of TOFA (Figure S1A) from 1 h p.i. onward, following removal of the virus inoculum. Total intracellular RNA and cell culture supernatant were collected at 16 h p.i. Interestingly, compared to the DMSO-treated control, no changes were observed in the levels of intracellular viral RNA (as detected by RT-qPCR) when infected Huh7 (Figure 2A) or MRC5 (Figure 2B) cells were treated with TOFA. However, by plaque assay we measured a 4-log reduction in infectious viral progeny released from Huh7 cells (Figure 2C) and an approximately 2.5-log reduction in the case of MRC5 cells (Figure 2D). When using TVB-2640, a small-molecule inhibitor that blocks the activity of FASN downstream of ACC (Figure 1A), similar trends were again observed in Huh7 cells, with the intracellular viral RNA levels remaining unaltered while the infectious progeny titers were reduced approximately 30-fold upon compound treatment (Figure 2E and 2F). Taken together, our data indicated that the DNL branch of lipid metabolism plays a proviral role in MERS-CoV replication.

**Figure 2.**
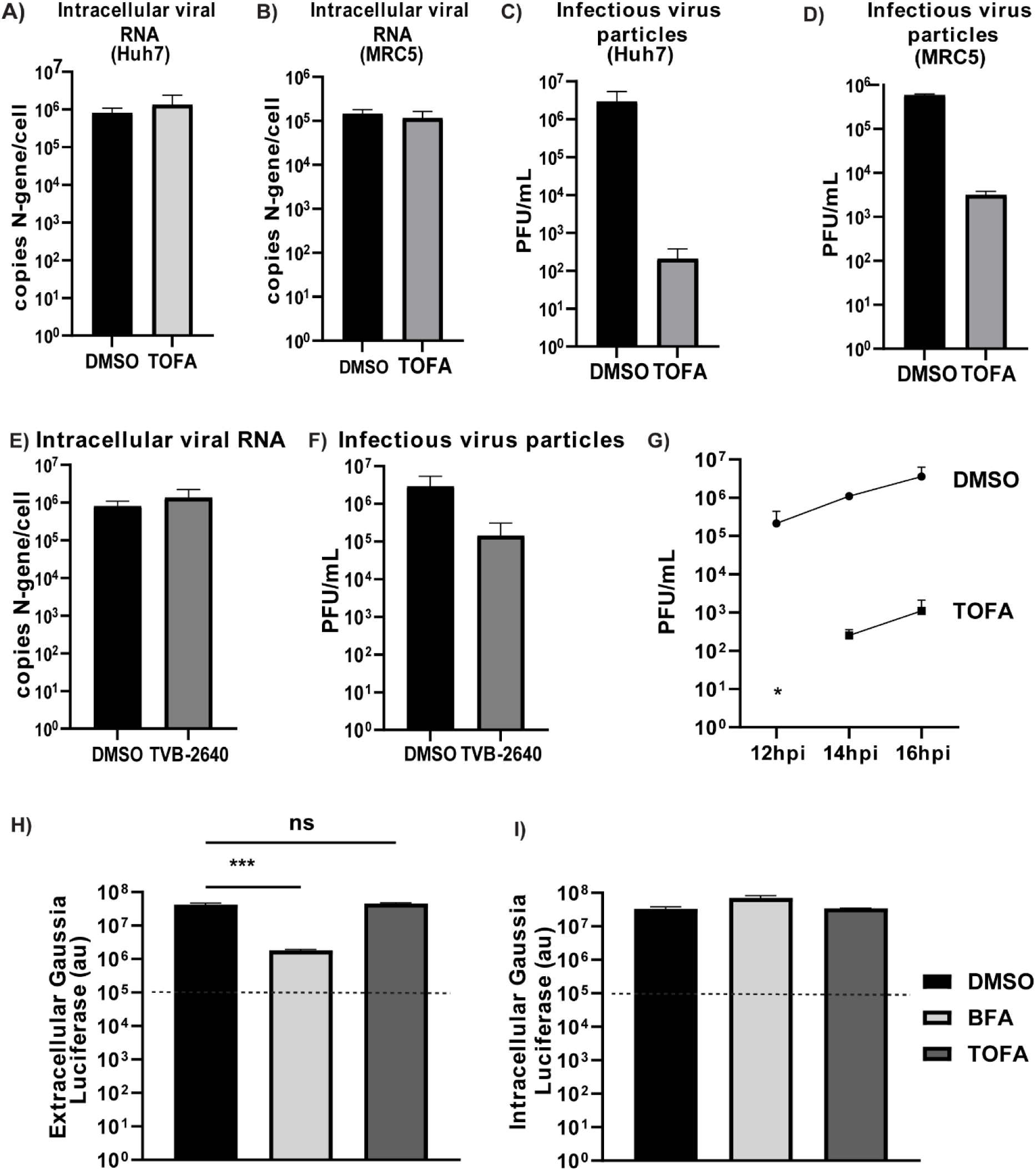
DNL inhibition strongly reduces the release of infectious MERS-CoV progeny, without affecting intracellular viral RNA synthesis. A) Huh7 cells or B) MRC5 were infected with MERS-CoV (MOI 5) and treated with either DMSO (vehicle control) or 10 μM TOFA from 1 h p.i. onward. Cell lysates were harvested at 12 h p.i. (Huh7 cells) or 16 h p.i. respectively, and intracellular viral RNA copies were measured by RT-qPCR. C) and D) Infectious viral progeny was quantified by plaque assay on Huh7 cells, using the harvested culture supernatant (see panel A and B, respectively). E) Huh7 cells were infected with MERS-CoV (MOI 5) and treated with either DMSO (vehicle control) or 15 μM TVB-2640 from 1 h p.i. onward. At 16 h p.i., cells were harvested, and intracellular viral RNA copies were measured by RT-qPCR. F) Infectious viral progeny from TVB-2640-treated and untreated Huh7 cells (see panel E) was quantified by plaque assay on Huh7 cells. G) Huh7 cells were infected with MERS-CoV (MOI 5) and treated with either DMSO (vehicle control) or 10 μM TOFA from 1 h p.i. onward. Samples were collected at 12, 14, and 16 h p.i. Infectious viral progeny was quantified by plaque assay on Huh7 cells. While at 12 h p.i the number of infectious virus particles was below the detection limit of the assay, virus titers increased over time at 14 h p.i and 16 h p.i. H) Analysis of the secretory pathway’s functionality upon TOFA treatment. Huh7 cells were transfected with a plasmid expressing *Gaussia* luciferase, a naturally secreted reporter protein. From 8 h post transfection., transfected cells were treated with either 10 μM TOFA or 5 μM Brefeldin A. At 24 h post compound addition, the levels of extracellular *Gaussia* luciferase secreted in the supernatant were measured using an enzymatic assay (see Materials and Methods). I) The cell monolayers from the same wells (panel H) were also lysed and intracellular *Gaussia* luciferase expression was measured and compared with the amount of secreted reporter protein. Dotted lines indicate the background levels measured. Data are represented as mean ± SD of two (D), three (B, E, G) or four (H, I) biological replicates. Statistical significance was calculated using one-way ANOVA and applying Bonferroni multiple comparison correction, *** p< 0.001

When MERS-CoV-infected Huh7 cells were treated with TOFA, the amount of infectious viral progeny was found to slowly increase from undetectable at 12 h p.i. to about 10^3^ PFU/ml by 16 h p.i. Although the viral titers showed an increase over time, a systematic 4-log difference was observed when comparing the vehicle control and the TOFA-treated samples (Figure 2G). Overall, these data led us to conclude that pharmacological inhibition of ACC, without affecting viral RNA synthesis, severely hampered a later step in the coronavirus replication cycle, such as virus assembly, egress, or virus particle infectivity.

Because of the significant reduction in extracellular infectious viral progeny following compound treatment, we sought to understand whether the pharmacological inhibition of ACC affected the host cell’s secretory pathway, which is involved in virion release. Using a *Gaussia* luciferase reporter assay (43), we observed that TOFA did not affect the integrity of the secretory pathway, while Brefeldin A, a well-characterized protein trafficking inhibitor, strongly inhibited the secretion of *Gaussia* luciferase in the supernatant (Figure 2H). There were no differences in the intracellular levels of *Gaussia* luciferase across all conditions (Figure 2I), which excluded a general effect of TOFA on protein synthesis. Taken together, our data pointed to an intact secretory pathway in the TOFA-treated cells, without a negative, off-target effect of the compound on the cellular translation machinery, suggesting that the reduction in extracellular infectious particles was not due to a virus egress defect.

### TOFA alters the host cell lipidomic profile and induces lipolysis

To gain insight into TOFA’s impact on host pathways that are relevant for virus assembly and release, uninfected cells were treated with the compound and harvested 12 h post treatment for targeted lipidomics analysis. This revealed that TOFA treatment induced profound changes in the host cell’s lipidomic profile, including increased levels of phosphatidylcholine, phosphatidylserine, lysophospholipids, and lipids of the ceramide biosynthetic pathway (Figure S2A). Strikingly, the levels of neutral lipids including triacylglycerols (TG), cholesterol esters (CE) and diacylglycerols (DG) were significantly reduced (Figure S2A), while free fatty acids (FFA) increased. These latter effects strongly suggested a metabolic switch in TOFA-treated cells towards the hydrolysis of triacylglycerols and diacylglycerols into their constituent molecules of acylglycerols and FFAs, a process also known as lipolysis. Lipolysis ultimately results in the degradation of LDs, which constitute the cellular neutral lipid storage compartments (44).

To corroborate the above findings, we used immunofluorescence microscopy and LD labeling with BODIPY 493/503, a well-known neutral lipid dye, and observed a reduced amount of LDs in TOFA-treated cells compared to DMSO-treated control cells (Figure S2B). Our results strongly suggested that pharmacological inhibition of ACC and thus FFA synthesis has a broader effect on lipid metabolism than previously described (42) and induces a compensatory shift toward lipolytic pathways. Similar effects were observed in MERS-CoV-infected cells in a time-dependent manner. At 9 h p.i. (8 h after the start of TOFA treatment), lipid droplets were still present at detectable levels in TOFA-treated infected and uninfected cells (Figure 3A and 3B, panels ii), albeit in somewhat reduced amounts in comparison to the DMSO-treated control cells (Figure 3A and 3B, panels i). However, by 12 h p.i. (11 h after the start of TOFA treatment), there was a prominent difference between the DMSO- and TOFA-treated cells (Figure 3A and 3B, compare panels iii and iv), with the abundance of LDs being strongly reduced in the latter. Taken together, TOFA treatment appeared to induce a gradual metabolic rewiring towards lipolytic pathways in both mock-infected and MERS-CoV-infected cells.

**Figure 3.**
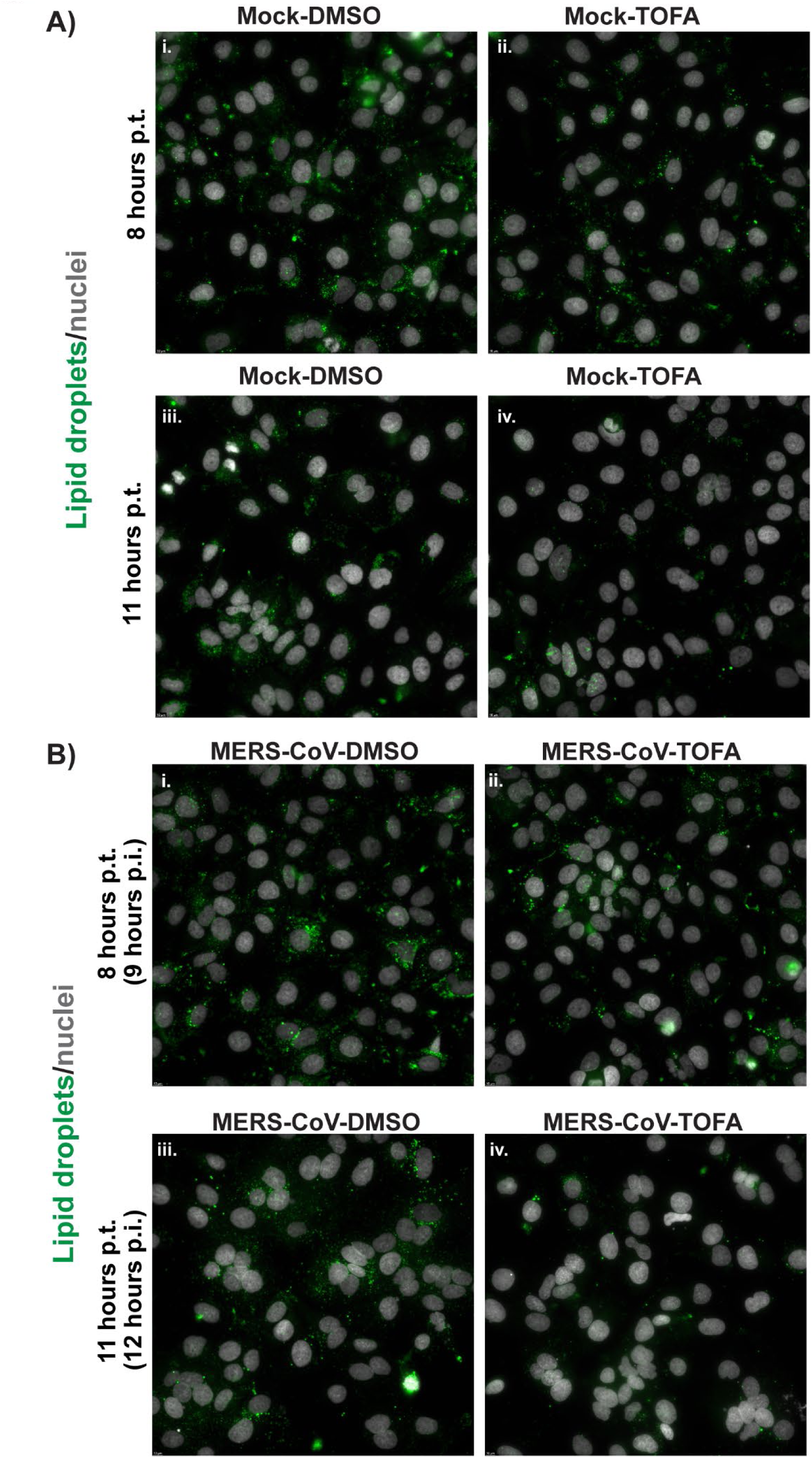
TOFA induces lipolysis in uninfected and MERS-CoV-infected cells. A) Uninfected Huh7 cells were treated with DMSO or 10 μM TOFA. B) Huh7 cells were infected with MERS-CoV (MOI 5) and treated with either DMSO (vehicle control) or 10 μM TOFA from 1 h p.i. onward. For both uninfected and infected cells, samples were collected at 8 and 11 h p.t. The next day the fixed cells were stained with BODIPY 493/503 (lipid droplets), and Hoechst 33342 (nuclei) and the presence of lipid droplets was analyzed by wide-field fluorescence microscopy.

### Subcellular localization of MERS-CoV structural proteins upon ACC inhibition

Next, we studied whether the trafficking of MERS-CoV envelope proteins was affected by ACC inhibition. To this end, we investigated their subcellular localization by immunofluorescence labeling and confocal microscopy, while using antibodies recognizing well-characterized host marker proteins to visualize specific compartments of the secretory pathway. Specifically, antibodies against S and M or S and E proteins were used for the simultaneous labeling of two of the envelope proteins, while in the same specimens antibodies against PDI, ERGIC-53, or giantin were used to stain the endoplasmic reticulum (ER), ER-Golgi intermediate compartment (ERGIC), and Golgi complex, respectively. As previously reported for other coronaviruses (26, 45, 46), in untreated infected cells, viral envelope proteins co-localized in both ERGIC (Figure S3) and Golgi membranes (Figure 4A-B). After ACC inhibition, limited changes in the intracellular co-localization patterns of the envelope proteins were observed. For example, in the DMSO-treated control cells, the S protein co-localized with the giantin marker in the Golgi complex (white arrows, Figures 4A and 4B, panel i, yellow spots). Post-TOFA treatment, this co-localization was not observed with the same intensity and the signal for both S protein and giantin seemed more dispersed (Figures 4A and 4B, panel ii). Furthermore, while in the untreated control cells E (Figure 4A) or M (Figure 4B) proteins co-localized with giantin (white arrows in Figures 4A and 4B, panel iii, yellow spots), these co-localizations were no longer apparent after TOFA treatment (Figures 4A and 4B, panel iv). In combination, TOFA treatment resulted in a reduced triple co-localization between S and E proteins with the giantin marker (Figure 4A) or S and M proteins with the giantin marker (Figure 4B) (panels v., white arrows, in comparison with panels vi.). Though the changes upon ACC inhibition were limited in nature, they did suggest that viral envelope protein trafficking to the Golgi complex was altered compared to that in untreated MERS-CoV-infected cells.

**Figure 4.**
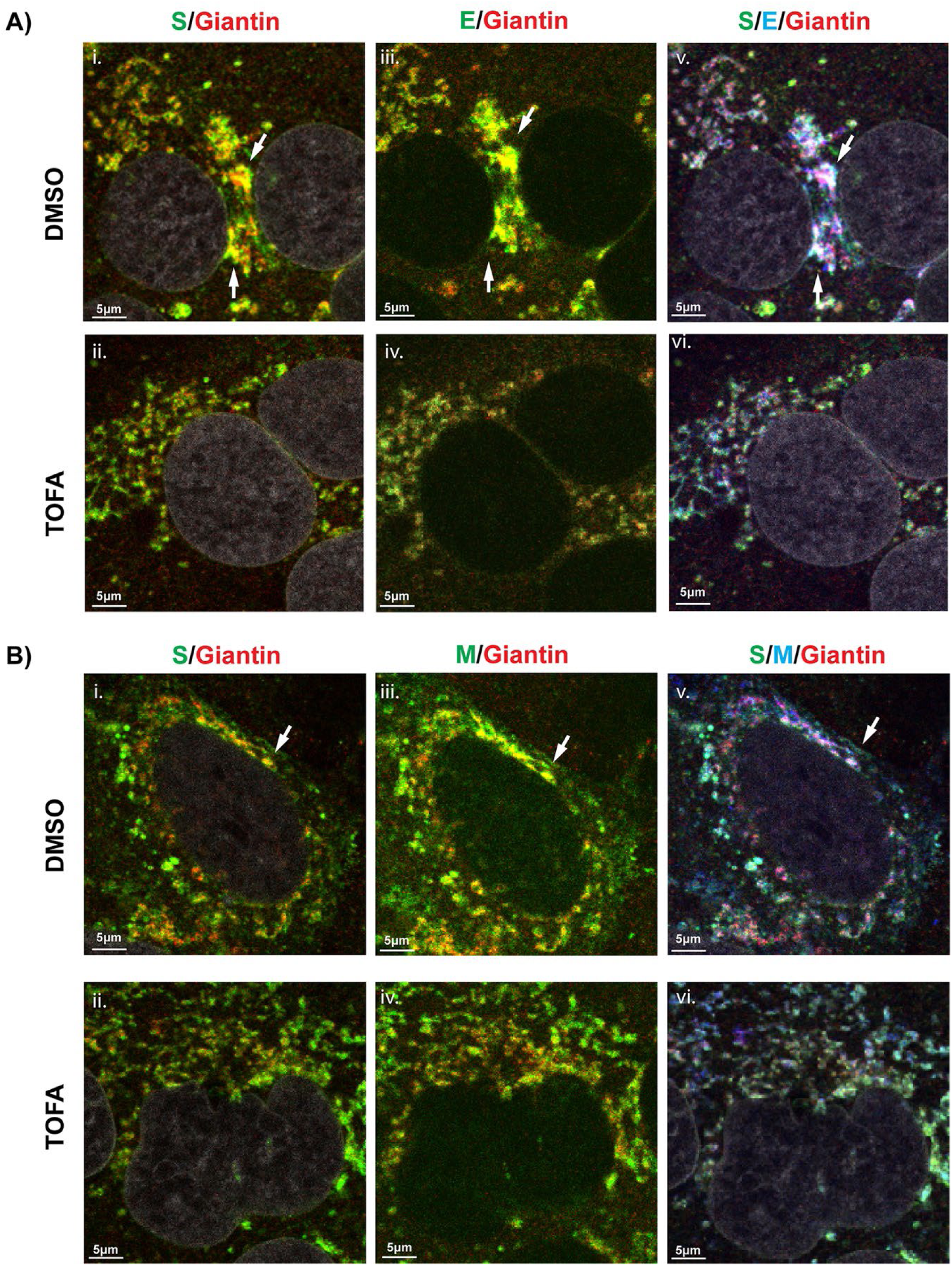
Subcellular localization of MERS-CoV envelope proteins following TOFA treatment. Huh7 cells were infected with MERS-CoV, treated with TOFA as previously described, fixed at 12 h p.i and analyzed using immunofluorescence microscopy. Cells were triple-labelled for A) S (green, panels i. and ii. ) and or E proteins (green, panels iii. and iv.) in combination with labeling for giantin (red), as a marker for the Golgi complex or B) S (green, panels i. and ii.) and M (green, panels iii. and iv.) proteins in combination with labeling for giantin (red). Confocal images are representative of at least two independent biological replicates. White arrows point to co-localization foci of two or three markers.

### TOFA treatment affects intracellular processing and trafficking of MERS-CoV envelope proteins

Using Western blot analysis, we further investigated the status of the MERS-CoV structural proteins S, E, M, and N in both extracellular samples (culture supernatants from infected and TOFA-treated cells) as well as cell lysates.

Only trace amounts of the N protein could be detected in the supernatant collected from TOFA-treated infected cells (Figure 5A), while the amounts of the three envelope proteins were below the detection limit of our assay. Importantly, this result excluded the possibility that the decrease in viral infectivity titers in supernatants from TOFA-treated cells (Figure 2C) was merely due to the production of non-infectious virus particles.

**Figure 5.**
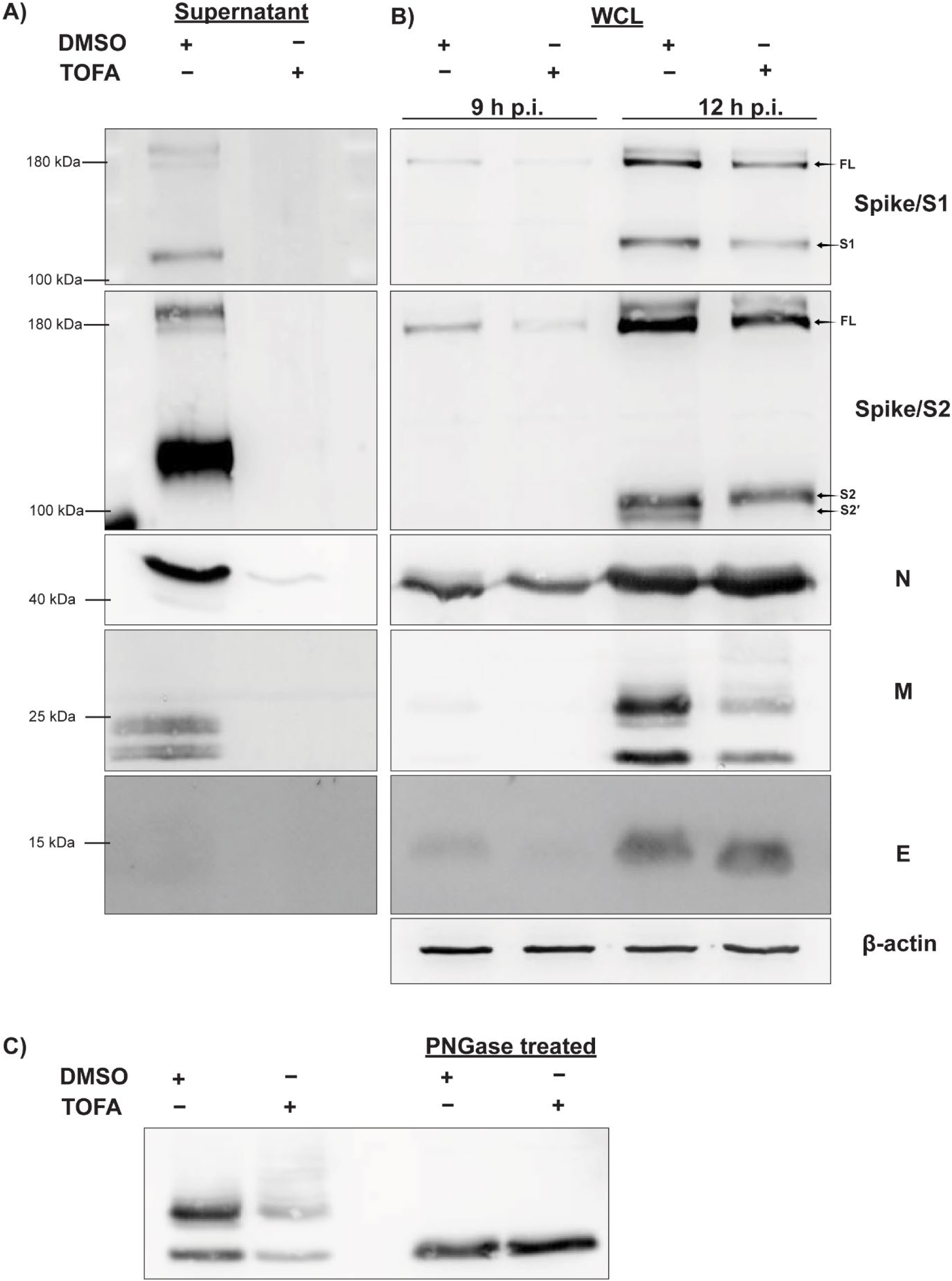
Effect of TOFA treatment on MERS-CoV structural protein expression and post-translational modifications. Expression levels of S protein (using separate antibodies against the S1 and S2 subunits), N protein, M protein, and E protein were analyzed by western blot, using β-actin as internal control. A) Huh7 cells were infected with MERS-CoV and treated with DMSO or TOFA as previously described. Supernatants were collected at 16 h p.i. and progeny virus particles were pelleted through a sucrose cushion by ultracentrifugation, lysed, and used for Western blot analysis. B) Huh7 cells were infected with MERS-CoV and DMSO- or TOFA-treated as previously described, and samples (WCL: Whole Cell Lysate) were collected at 9 and 12 h p.i. for analysis by Western blotting. C) Western blot for PNGase assay (described in Materials and Methods) using lysates collected from MERS-CoV-infected Huh7 cells and collected at 12 h p.i. Blots are representative of at least two independent biological replicates.

Intracellularly, only the overall levels of N protein remained more or less unaffected at 9 and 12 h p.i. in the presence of TOFA. Decreased levels of full-length (FL) S glycoprotein as well as its S1 and S2 fragments were observed, at 9 h and 12 h p.i. Moreover, after TOFA treatment, we could not detect a second band below the S2 fragment, which likely corresponds to the S2′ fragment (47), suggesting a decreased transport of the S protein to the trans-Golgi network where the furin protease responsible for this cleavage is located. We also observed that the E protein consistently migrated slightly faster after compound treatment, again suggesting changes in the protein’s post-translational modifications (Figure 5B).

The M protein was detectable as the usual double band, indicative of the two different glycosylation states of M (48), migrating around 15-25 kDa (Figure 5B). However, following compound treatment, the blot for the M protein also revealed a smear of products with a higher molecular weight (in the 20-35 kDa range), suggesting a change in the post-translational maturation and/or trafficking of the M protein. The smear disappeared after glycosidase treatment of the cell lysates with PNGase (Figure 5C), confirming that it resulted from a change in the glycosylation status of M, which was likely linked to a disruption of the protein’s intracellular trafficking.

### Electron microscopy reveals that TOFA treatment interferes with MERS-CoV assembly and release

Overall, our data confirmed that TOFA treatment caused a major defect in the release of infectious MERS-CoV progeny, and excluded the possibility that large amounts of non-infectious particles were secreted instead. Our subsequent microscopy and biochemical studies pointed towards changes in the post-translational modification and trafficking of all three MERS-CoV envelope proteins, which might disrupt the normal intracellular assembly of new virions.

We employed transmission electron microscopy (TEM) to directly visualize the impact of TOFA treatment on the viral replication cycle. Huh7 cells were infected with MERS-CoV at MOI 5 and treated with the compound from 1 h p.i. onward. Cells were fixed at 12 h p.i. and analyzed with TEM. We observed that pharmacological inhibition of ACC did not affect the formation or morphology of virus-induced ROs (DMVs and CMs) in comparison to the vehicle-treated control cells (Figure 6A, panels i and ii), which aligns with our previous data regarding the unaffected intracellular viral RNA levels upon TOFA treatment (Figure 2A). Moreover, while extracellular progeny virions were abundantly found at the plasma membrane of untreated MERS-CoV-infected cells, they were hardly present in the TOFA-treated samples (Figure 6B, panels i. and ii.), in agreement with the strong reduction in progeny virus titers observed when ACC is inhibited (Figure 2B).

**Figure 6.**
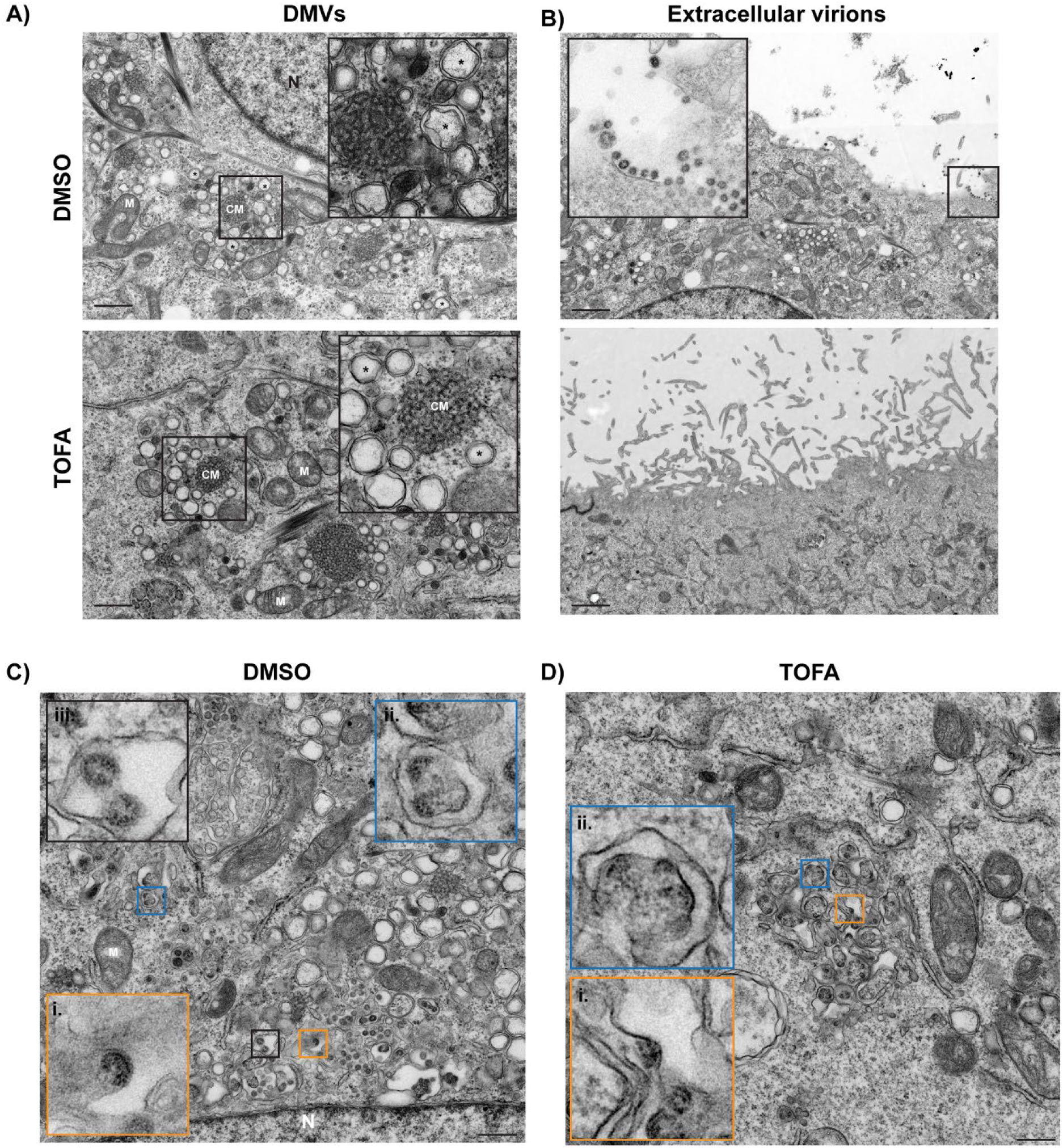
TEM analysis of TOFA-treated Huh7 cells infected with MERS-CoV. Cells were infected with MOI 5, treated with TOFA or with DMSO and fixed and processed for electron microscopy at 12 h p.i. A) Morphologically comparable virus-induced DMVs (asterisks) can be readily observed in both conditions. Close-ups of the boxed areas are provided in the insets. B) Extracellular virions abundantly present in the extracellular space of DMSO-treated MERS-CoV-infected cells (i, inset: zoomed-in image of the boxed area), but not in the extracellular space of TOFA-treated and virus-infected cells (ii). C) Consecutive stages of MERS-CoV particle assembly in DMSO-treated and infected cells; assembly events were split in two categories for simplicity (see Materials and Methods), denoted as early-stage assembly events of budding nucleocapsid complexes in membranes of the secretory pathway (panels i and ii) and late-stage assembly events including nearly fully-formed virus particles and virions fully-budded inside large vesicles (iii). D) In TOFA-treated MERS-CoV-infected cells early assembly stage events, as seen in panels (i) and (ii), were abnormally abundant, revealing a stalling of virus particle assembly in the presence of the compound. Mitochondria are annotated with M, convoluted membranes with CM, and cell nuclei with N. Scale bar is 200 nm.

Our ultrastructural studies further revealed that the consequences of TOFA treatment hindered virus particle assembly. In untreated control samples (Figure 6C), different stages of virion budding could be detected, from early stages in which the nucleocapsid starts to be engulfed by a curved membrane (Figure 6C, panel i and ii) to (nearly) fully budded virions (panel iii), with the latter representing most virions. However, in TOFA-treated samples, the early stages of virus particle assembly were particularly abundant (Figure 6D), with exceptionally large areas containing particles representing this stage (Figure 6D, panels i and ii). This strongly suggested that TOFA treatment interferes with the normal progress towards a fully budded virus particle, leading to an unusual accumulation of nucleocapsids in an initial budding state.

### Inhibition of palmitoyltransferase enzymes mimics the effects of ACC inhibition

One of the many end products of the DNL pathway is the fatty acid palmitate. Palmitate, or palmitic acid, plays an important role in various cellular processes: as a signaling molecule, a structural element of various lipids, and by participating in posttranslational modifications, such as protein palmitoylation catalyzed by palmitoyltransferase enzymes (49). Coronavirus S and E proteins contain a conserved C-terminal cysteine-rich region next to their transmembrane domain, which can serve as a palmitoylation substrate (Figure 7A) (50). Previous studies in cells infected with the murine hepatitis (MHV), transmissible gastroenteritis (TGEV), SARS-CoV, and SARS-CoV-2 coronaviruses, have shown that their S proteins are post-translationally palmitoylated (51–58), while the cysteine-rich regions of the E proteins of SARS-CoV, MHV, and infectious bronchitis virus (IBV) were also found to be palmitoylated (59–62). Since there is no experimental data on the palmitoylation status of the MERS-CoV S and E proteins and the potential role of this posttranslational modification in assembly, we initially tested the importance of palmitoyltransferase enzymes for MERS-CoV replication using 2-bromopalmitic acid (2-BP), a small-molecule inhibitor with a broad effect on palmitoyltransferase enzymes (Figure 1A) (63). In an earlier antiviral compound screen (6), 2-BP was shown to inhibit MERS-CoV replication in a multi-cycle experimental setup, but its mechanism of action was not explored. We now employed a single-cycle setup as described above: Huh7 cells were infected with MOI 5, treated with 2-BP from 1 h p.i. onward, and at 16 h p.i. culture supernatants and total intracellular RNA were collected. Interestingly, we observed that inhibition of palmitoyltransferase enzymes induced similar changes as the ACC inhibition (Figure 7B and 7C). Compared to the untreated control, no major changes were observed in the levels of intracellular viral RNA detected by RT-qPCR in cells treated with 2-BP (Figure 7B). However, there was a ∼250-fold reduction in the extracellular infectious progeny titers (Figure 7C), aligning with the hypothesis that limited availability of palmitate after DNL inhibition may be partly responsible for the observed assembly impairment.

**Figure 7.**
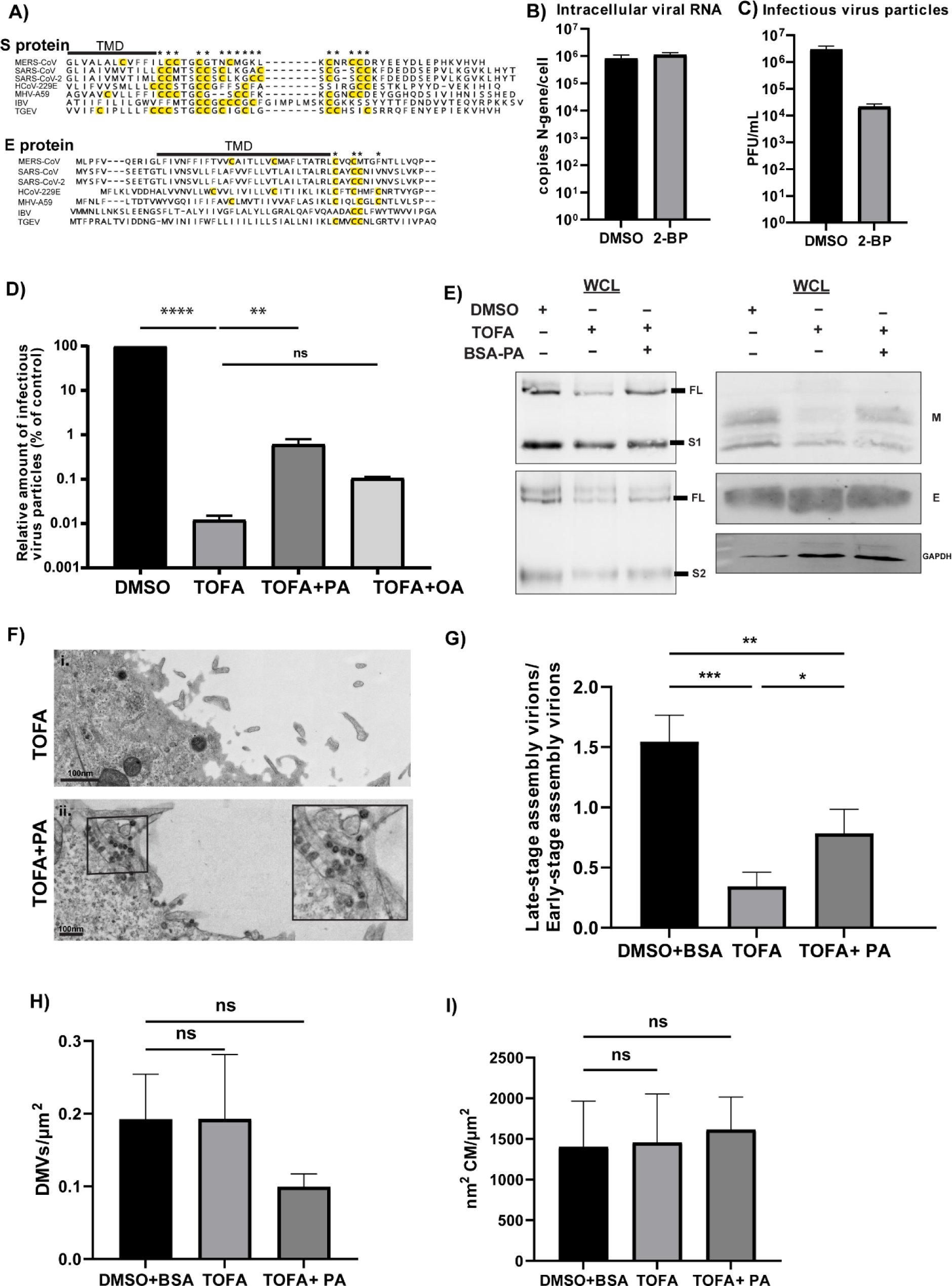
Exogenous palmitic acid supplementation partially rescues MERS-CoV assembly after TOFA treatment. A) Alignment of S and E protein C-terminal sequences from selected coronaviruses using the following accession numbers: MERS-CoV (NC_019843), SARS-CoV (NC_004718) SARS-CoV-2 (NC_045512), MHV (AY700211), TGEV (AJ271965), IBV (NC_001451), HCoV-229E (NC_001451). Cysteines next to the transmembrane domain that can serve as palmitoylation substrates are highlighted in yellow and noted with an asterisk. B) Huh7 cells were infected with MERS-CoV (MOI 5) and treated with either DMSO (vehicle control) or 30 μM 2-BP from 1 h p.i. onward. At 16 h p.i., cell lysates were harvested, and intracellular viral RNA copies were measured by RT-qPCR. C) Infectious viral progeny was quantified by plaque assay on Huh7 cells. Data are represented as mean ± SD of three independent experiments. D) Huh7 cells were infected with MERS-CoV (MOI 5) and treated from 1 h p.i. onward with DMSO supplemented with 100 μM BSA, TOFA, TOFA supplemented with 100 μM BSA-PA, or TOFA supplemented with 100 μM BSA-OA. At 16 h p.i., samples were collected, and infectious viral progeny was quantified by plaque assay on Huh7 cells. Data are represented as mean ± SD of two independent experiments. Statistical significance was calculated using one-way ANOVA and applying Dunnet multiple comparison correction, **** p< 0.0001 and ** p< 0.01, ns: not significant. E) Huh7 cells were infected with MERS-CoV and treated with DMSO, TOFA or TOFA with BSA-PA from 1 h p.i. onward as previously described, and samples were collected at 12 h p.i. for analysis by western blotting. Blots are representative of at least two independent biological replicates. F-I) Huh7 cells were infected with MERS-CoV and treated as described above. At 12 h p.i., samples were fixed and processed for TEM. F) Post addition of palmitic acid, the number of extracellular virus particles present in TOFA-treated virus-infected cells is increased. Scale bar is 100nm. G) Virions either in the late-stage (advanced/ fully assembled particles) or early-stage were quantified (see Materials and Methods), and the ratio of both classes was determined. H) Number of DMVs per µm^2^ of cytoplasm observed (see Materials and Methods) in the DMSO and BSA-treated, TOFA-treated or TOFA and BSA-PA-treated virus-infected cells. I) In the same samples, the area of CMs (in nm^2^) was also determined per µm^2^ of cytoplasm (see Materials and Methods). Data are represented as mean ± SD of three separate stiches. Statistical significance was calculated using one-way ANOVA and applying Tukey multiple comparison correction, *** p< 0.001, ** p< 0.01, and * p<0.05.

### Addition of exogenous palmitic acid partially rescues assembly and release of MERS-CoV progeny

Based on the possible involvement of palmitoylation defects in the observed disturbances of MERS-CoV assembly, we investigated whether assembly could be restored by supplementing some of the end products of the DNL pathway, such as palmitic acid or oleic acid. To this end, Huh7 cells were infected with MERS-CoV at MOI 5 and from 1 h p.i. onward they were treated with 10 μM TOFA or TOFA supplemented with 100 μM BSA-conjugated palmitic acid or BSA-conjugated oleic acid. A treatment with 100 μM BSA without conjugated fatty acid served as a vehicle control for the fatty acids (Figure 7D). Fatty acid concentrations where chosen based on previous cytotoxicity assays, using concentrations that would not reduce cell viability below 90% (Figures S1E and S1F), thus aiming to avoid potential confounding effects of lipotoxicity-induced cell death. Interestingly, we observed that the addition of exogenous palmitic acid conjugated with BSA partially restored the release of infectious MERS-CoV progeny in the supernatant (by almost 100-fold), while addition of BSA-conjugated oleic acid or BSA alone did not (Figure 7D).

Moreover, the effects of TOFA treatment that we had observed at the level of individual envelope proteins (Figure 5) could be partially reversed by addition of palmitic acid, resulting in intracellular protein levels and post-translational modifications that were closer to those seen in samples from untreated infected cells (Figure 7E). Electron microscopy analysis of large EM data sets also revealed an increase in the number of extracellular virions after palmitic acid addition (Figure 7F), while intracellularly there was a significant decrease in the proportion of virions arrested in the early assembly stage (Figure 7G, with representative images provided in Figure S4). In contrast, a quantitative analysis of the abundance of DMVs (Figure 7H) and CMs (Figure 7I) did not reveal significant changes in comparison to the DMSO-treated control, neither upon TOFA treatment nor upon addition of palmitic acid. Overall, our data indicate an important role for palmitic acid during MERS-CoV assembly, potentially involving palmitoylation of the S and E structural proteins.

## Discussion

Being enveloped positive-strand RNA viruses, coronaviruses depend on the host cell’s endomembrane system to facilitate and support their propagation in several ways. They transform intracellular, ER-derived membranes into the ROs that accommodate viral RNA synthesis and hijack membranes of the secretory pathway to generate enveloped virus particles. Due to this extensive involvement of host membranes, lipid metabolism pathways have been implicated in coronavirus replication (5, 6). After the emergence of SARS-CoV-2, several studies touched upon the potential reprogramming of lipid metabolism in the context of coronaviral replication (8, 10, 11, 39), but few mechanistic details were resolved. Here, we report on the importance of lipid homeostasis in MERS-CoV-infected cells for viral propagation to take place, as perturbations of the fatty acid biosynthesis pathway were found to have a strong inhibitory effect, specifically on the assembly and/or release of viral progeny.

Due to the extensive remodeling of the ER during coronavirus RO biogenesis, it has been speculated that fatty acid biosynthetic pathways would be activated to fuel the formation of these membrane structures (6, 10). However, our data show that perturbation of DNL does not affect MERS-CoV RNA synthesis or RO biogenesis (Figure 2A, 2B, 2E and 6A), indicating that – contrary to previous speculations - DNL is not among the host cell pathways critically required for coronaviral RO formation (6, 10). Our data are in partial agreement with published literature on SARS-CoV-2. Earlier studies have shown that lipid metabolism pathways, especially anabolic branches such as DNL and not catabolism of fatty acids (like β-oxidation), play a general proviral role in coronavirus replication (6, 10, 39), which is also corroborated by our data when using inhibitors of ACC and FASN.

A previous SARS-CoV-2 study (10), using TOFA among other small-molecule inhibitors of the DNL pathway, showed that DNL inhibition impedes SARS-CoV-2 replication. However, using immunofluorescence microscopy, Williams et al, observed a change in the intracellular distribution of double-stranded RNA (dsRNA) foci, which was interpreted to represent a change in the formation of viral ROs. In our study with MERS-CoV, such a dsRNA pattern change was not observed (Figure S5), while electron microscopy analyses of large EM data sets showed no alterations in the morphology (Figure 6) or abundance (Figure 7H-I) of viral ROs. These differences could be attributed to technical details of the immunolabeling protocol or the use of different coronavirus species and host cells. Another technical difference was the use of a single-cycle of viral replication setup in our study, in contrast to the multi-cycle approach and longer experimental timeframes applied in most of the previously published studies (6, 10, 39, 64). By using a single-cycle approach, we aimed to synchronize the different steps of the viral cycle (entry, RO formation, peak of viral RNA and protein synthesis, particle assembly) and study them at defined timepoints.

Our data indicate that the DNL pathway, and in particular palmitate, one of its end products, plays a role in MERS-CoV particle assembly or maturation (Figure 7) rather than in RO formation. Palmitoylation of the S and E proteins has been reported to be important for the replication of other coronaviruses, like MHV, SARS-CoV, and SARS-CoV-2 (51–62), making it tempting to assume a similar involvement for MERS-CoV. However, palmitoylation of MERS-CoV structural proteins has not been investigated directly, in infected cells or overexpression/virus-like-particle systems. Consequently, the number of palmitoylated cysteines, and their role and position with respect to the transmembrane domain of the MERS-CoV S and E proteins remain unknown.

Depending on the coronavirus studied, disruption of palmitoylation seems to have divergent effects on the replication cycle. For example, palmitoylation of the MHV S protein was reported to be necessary for its interaction with the M protein and assembly of virus particles (51), while palmitoylation of the SARS-CoV or SARS-CoV-2 S protein is not required for its interaction with the M protein (53, 55). It is unknown and rather difficult to predict what the importance of palmitoylation of the MERS-CoV envelope proteins might be, since the molecular interactions between coronaviral S and M protein are generally not well defined. In the coronaviruses studied so far, palmitoylation of S and E proteins seems to increase protein stability and to decrease their turnover rate (61, 62). This could also be the case for MERS-CoV, as suggested by the decrease we observed in intracellular S protein levels after ACC inhibition (Figure 5B). Attempts to use a triple-mutated MHV E protein lacking all potentially palmitoylated cysteines to generate virus-like particles (VLPs) or recombinant mutant MHV, resulted in particles incapable of proper assembly and release in both cases (61, 62). Interestingly, this triple-mutated MHV E protein also exhibited faster migration during SDS-PAGE (61), which is in line with our western blot observations after ACC inhibition (Figure 5B) and the partial recovery of the protein’s normal migration after addition of palmitic acid (Figure 7E). Besides impacting their stability, another common effect of palmitoylation on the S and E proteins is the influence on their affinity for specific membrane lipid nanodomains (55, 61). Lack of S and E protein palmitoylation might lead to their association with membranes of different lipid composition than what is necessary for efficient assembly, eventually leading to disruption of the regular assembly events. This is partially supported by our immunofluorescence microscopy data, since we observed reduced co-localization of the S and E proteins with a Golgi compartment marker (Figure 4).

The most abundant viral envelope protein and a key driver of particle assembly is the M protein, which has not been reported to be palmitoylated, but it is known to be glycosylated on its short N-terminal ectodomain. Upon TOFA treatment, the M protein exhibited differences in its glycosylation pattern (Figure 5B and 5C). Being on the luminal side of the membrane, the glycans decorating the MHV or SARS-CoV M proteins do not seem to have a crucial role in virion assembly and release, nor in M protein interactions with the S and E proteins (65, 66). However, these previous studies have mainly explored the use of mutated M proteins that lacked the specific amino acid residues normally used for glycan conjugation. In our study, following TOFA treatment, we observed M proteins carrying longer, possibly untrimmed, glycan chains. Trimming of glycan chains takes place as proteins progress from the ER to the Golgi complex, as part of the ER’s quality control machinery for misfolded proteins. Glycans are further processed, trimmed, and extended during protein trafficking through the Golgi complex, as part of normal glycoprotein maturation (67, 68). Thus, the M protein’s glycosylation differences observed in our study could signal trafficking issues that lead to delayed or aberrant processing of its (predicted) single N-linked glycan. Although these changes may not affect virus assembly, they are in line with our immunofluorescence microscopy observations regarding the M protein’s reduced colocalization with giantin (Figure 4). According to the western blot results (Figure 5C), about half of the total amount of M protein still appears to be properly processed and trafficked, but the impact of the reduced or delayed availability of M protein at the site of virus assembly is difficult to predict.

In conclusion, the observed major impairment of the assembly and release of infectious MERS-CoV progeny upon ACC inhibition may well be a multifactorial phenomenon. In our TEM images (Figure 6D), inhibition of the fatty acid synthesis pathway resulted in a striking increase of the number of early assembly/budding events relative to (nearly) fully-formed virus particles. TOFA treatment may affect virus particle assembly by reducing palmitoylation of S and E proteins, which can potentially result in their instability and increased turnover. Simultaneous changes in M protein trafficking may contribute to an overall imbalance in the viral envelope proteins needed for interactions with each other, host lipids, and viral nucleocapsids, ultimately affecting the pace and quality of virion assembly and budding. In addition to its presumed impact on the trafficking and post-translational modification of viral envelope proteins, TOFA treatment may also affect the properties of unidentified host factors that are directly or indirectly involved in coronavirus assembly. This could also contribute to the general disruption of lipid metabolism pathways and the metabolic switch towards lipolytic pathways that we observed in both mock- and MERS-CoV-infected cells (Figure 3 and S3), which could contribute to an unfavorable microenvironment and host membrane composition for efficient virion assembly. The proviral role of *de novo* lipogenesis in MERS-CoV replication also highlights its potential as a target for the development of host-directed antiviral therapeutics against coronaviruses.

## Materials and Methods

### Cells

Huh7 cells (kindly provided by Dr. Ralf Bartenschlager, Heidelberg University, Germany) and MRC5 cells (purchased from ATCC, CCL-171) were cultured in Dulbecco’s Modified Eagle Medium (DMEM) (Lonza) supplemented with 8% (v/v) fetal calf serum (FCS), 100 IU/mL penicillin/streptomycin, 2 mM L-glutamine, and non-essential amino acids, at 37°C and 5% CO2.

### Viruses and virus purification

MERS-CoV strain EMC/2012 (NC_019843) (69, 70) was kindly provided by Dr. Ron Fouchier, Erasmus Medical Center Rotterdam, the Netherlands, and working stocks were prepared using Huh7 cells. All work with live MERS-CoV was performed inside a biosafety level 3 facility at Leiden University Medical Center. Virus titrations by plaque assay were performed as described before (71). Huh7 cells were seeded in tissue culture-treated multiwell plates and the next day were infected with serial dilutions of MERS-CoV containing supernatant for 1 h at 37°C. The virus inoculum was removed and replaced by overlay medium containing 1.2% Avicel RC-581 (Cas. No.: 9004-34-6, FMC/IMCD Benelux) in DMEM supplemented with 2% FCS and 100 IU/mL penicillin/streptomycin. Cells were incubated for 3 days at 37°C, after which they were fixed with formaldehyde and stained with crystal violet.

Virus concentration by ultracentrifuge pelleting through a sucrose cushion (Figure 5A) Infected cell culture supernatants were harvested 16 h p.i. and clarified from cell debris by low-speed centrifugation at 1,200 g for 5 min at 4°C. The virus particles were pelleted through a 20% (w/v) sucrose cushion in TES buffer (0.02 M Tris-HCl, 1mM EDTA, 0.1 M NaCl, pH 7.4) by centrifugation in an SW41 Ti rotor (Beckman Coulter) for 3.5 h at 30,000 rpm and 4°C. Virus pellets were resuspended overnight in phosphate-buffered saline (PBS) at 4°C, lysed in 100mM Tris-HCl, pH 7.6 and 4% (w/v) sodium dodecyl sulfate (SDS), and used for SDS-PAGE and Western blot analysis.

### Small-molecule inhibitors

TOFA (5-tetradecyloxy-2-furoic-acid, cat. No.: HY-101068) and TVB-2640 (Denifanstat, cat. No.: HY-112829) were purchased from MedChemExpress. 2-bromopalmitic acid (2-bromohexadecanoic acid, cat. No: 21604) was purchased from Sigma. Compounds were dissolved in anhydrous DMSO for stock preparation. BSA-conjugated palmitate saturated fatty acid complex (PA) (cat. No.: 29558), BSA-conjugated oleate monounsaturated fatty acid complex (OA) (cat. No. 29557) and BSA control for BSA-fatty acid complexes (cat. No. 29556) were purchased from Cayman Chemicals.

### Cytotoxicity assays

Cell viability was monitored using CellTiter 96 aqueous non-radioactive cell proliferation reagent (Promega), following the manufacturer’s instructions. Cells were seeded in 96-well plates and incubated overnight at 37°C and 5% CO2. On the next day, the cells were treated with serial dilutions of the different compounds in DMEM containing 2% FCS, 100 IU/mL penicillin/streptomycin, 2 mM L-glutamine and non-essential amino acids. Cells were incubated for 22 h, and then 20 ml/well of MTS solution (3-(4,5-dimethylthiazol-2-yl)-5-(3-carboxymethoxyphenyl)-2-(4-sulfophenyl)- 2H-tetrazolium) were added. The cells were incubated for 2 h at 37°C, after which the absorbance was measured at 495 nm using the Envision multiplate reader (PerkinElmer). Cell viability was normalized against the DMSO-treated control.

### Quantitative PCR

Viral RNA in MERS-CoV-infected Huh7 cells was quantified by using RT-qPCR. To this end, samples were collected in Tripure Isolation Reagent (Sigma), intracellular RNA was isolated following the manufacturer’s instructions, and RT-qPCR was performed using the Applied Biosystems™ TaqMan Fast™ Virus 1-step Multiplex Master Mix for qPCR (ThermoFisher) with MERS-CoV-specific primers and probes (designed in-house as described before (72) ) in a CFX384 Touch RT-PCR detection system (Biorad). For absolute quantification, a standard curve was generated using 10-fold serial dilutions of an *in vitro* transcript containing the target sequences, generated using T7 RNA polymerase. The expression of human PGK1 gene (ThermoFisher, TaqMan gene expression Assay VIC-MGB, cat. No. #4448490, assay ID Hs00943178_g1) was used as an internal standard to ensure equal sample loading between conditions.

Gene expression analysis for host factors was performed using real-time RT-qPCR with iTaq SYBR Green Supermix (Biorad) in a CFX384 Touch RT-PCR detection system (Biorad). Primer sequences for *ACLY, ACC, FASN, SCD-1, DGAT1, ADRP, CPT1a, ACOX* and human β-actin were described elsewhere (6, 73). Quantification was performed by normalizing against the amount of human β-actin mRNA. Fold gene expression changes in MERS-CoV-infected cells relative to mock-infected cells were calculated by using the delta-delta Ct formula (2^–ΔΔCt^).

All oligonucleotide sequences and sources are provided in the Supplementary Table 1.

### *Gaussia* luciferase reporter assay

For monitoring the effect of TOFA on the secretory pathway’s activity, Pierce™ *Gaussia* luciferase glow assay kit (ThermoFisher, cat. No.: 16160) was used following the manufacturer’s instructions. Huh7 cells were seeded in 96-well plates and the next day, they were transfected with 150 ng of a Gaussia luciferase expression plasmid using Lipofectamine 3000 (ThermoFisher) at a 1:3 ratio of DNA to Lipofectamine. At 24 h post transfection, the cell culture supernatant was aspirated, and the cells were treated for 8 h with DMSO, TOFA, or brefeldin A. After incubation with the compounds, the cell culture supernatant containing secreted *Gaussia* luciferase was collected, and the cell monolayer was lysed for detection of intracellular luciferase. The luciferase signal output was measured at 485 nm using an Envision multiplate reader (PerkinElmer).

### Immunofluorescence microscopy

Huh7 cells were seeded on glass coverslips and left to attach overnight at 37°C and 5% CO2. The next day, cells were either mock- or MERS-CoV infected for 9 h or 12 h and then fixed with 3% paraformaldehyde. Immunofluorescence labelling was performed as described before (74). In this study, the following antibodies were used for immunofluorescence analyses: human monoclonal antibody against MERS-CoV Spike protein (kindly provided by Dr. Berend Jan Bosch, Utrecht University) (75), rabbit polyclonal anti-MERS-CoV M protein antibody (20, 76), rabbit polyclonal anti-MERS-CoV E protein antibody (ThermoFisher, cat. No.: PA5-143457), rabbit polyclonal anti MERS-CoV N protein antibody (Sino Biological, cat. No.: 40068-RP02), mouse monoclonal anti-dsRNA J2 antibody (Scicons), mouse monoclonal anti-PDI antibody (Enzo Life Sciences, ADI-SPA-891), mouse monoclonal ERGIC-53 G1/93 Ab (Enzo Life Sciences/ Alexis Biochemicals, ALX-804-602), mouse monoclonal anti-giantin antibody (Enzo Life Sciences/Alexis Biochemicals, ALX-804-600), BODIPY493/503 (ThermoFisher, cat. No.: D3922). For viral M and E protein labeling, an Alexa-647-conjugated donkey anti-rabbit secondary antibody (Jackson Laboratories, cat. No.: 711-605-152) was used, the S protein labelling was visualized with an Alexa-488-conjugated goat anti-human antibody (ThermoFisher, cat. No.: A11013), while for the N protein labeling an Alexa-488-conjugated goat anti-rabbit secondary antibody (ThermoFisher, cat. No.: A11008) was used. Cellular organelle markers and dsRNA were labelled using a Cy3-conjugated donkey anti-mouse antibody (Jackson Laboratories, cat. No.: 715-165-150). Leica Sp8 confocal and DM6 fluorescence microscopes were used for image acquisition. During confocal image acquisition, laser strength and gain were kept consistent between all samples and conditions. To obtain high-resolution images, an acquisition format of 1024x1024 pixels and speed of 400 Hz were selected and kept consistent between samples. Image acquisition settings were chosen to maximally reduce false-positive co-localization signal due to overexposure of the samples.

For image processing, Leica Application Suite X version 3.7 software was used. All images were processed in the same manner to increase brightness and contrast for this publication. Raw, unprocessed images are available upon request.

### Targeted lipidomics analysis

Comprehensive, quantitative shotgun lipidomics was carried out as described in detail elsewhere (77, 78). Briefly, samples were spiked with deuterated internal standards and lipids were extracted using methyl tert-butyl ether. The combined organic extracts were subsequently dried under a gentle stream of nitrogen and the dried extracts were dissolved in methanol:chloroform 1:1 containing 10 mM ammonium acetate. Lipids were then analyzed with a flow injection method at a flow rate of 8 µL/min applying differential ion mobility for lipid class separation and subsequent multiple reaction monitoring in positive and negative electrospray ionization mode. Using the Shotgun Lipidomics Assistant (SLA) software individual lipid concentrations were calculated after correction for sample input and their respective internal standards.

### Western blot and PNGase assay

After lysis of cells or purified virus particles, obtained as described above, the protein content of samples was quantified using the Pierce BCA protein assay kit (ThermoFisher). Samples were mixed with 4x Laemmli sample buffer, incubated at 96°C for 5 min, and separated using SDS-polyacrylamide gel electrophoresis (SDS-PAGE). Post separation, the proteins were transferred to Amersham Hybond-LFP 0.45 μm PVDF blotting membrane (Cytiva) using the semidry Trans-Blot Turbo Transfer System (Biorad). For the E protein, Amersham Hybond-LFP 0.2 μm PVDF blotting membrane was used instead. The membranes were blocked for 1 h at room temperature. For ACC and FASN, 5% bovine serum albumin (BSA) in TBS with 0.05% Tween-20 (TBST) was used as a blocking buffer, while for viral proteins 1% casein in PBS with 0.05% Tween-20 (PBST) was used instead. Membranes were incubated with primary antibodies overnight in 2.5% BSA in TBST or 0.5% casein in PBST respectively.

Antibodies used in this study were: ACC (Cell Signaling Technology, cat.no.: #3662), phospho-ACC (Ser79) (Cell Signaling Technology, cat.no.: #3661), FASN (Cell Signaling Technology, cat.no.: #3180), rabbit polyclonal anti-MERS-CoV S1 antibody (ThermoFisher, cat. No.: PA5-119581), rabbit polyclonal anti-MERS-CoV S2 antibody (ThermoFisher, cat. No.: PA5-81788), rabbit polyclonal anti-MERS-CoV M protein antibody (20, 76), rabbit polyclonal anti-MERS-CoV E protein antibody (ThermoFisher, cat. No.: PA5-143457), rabbit polyclonal anti-MERS-CoV N protein antibody (Sino Biological, cat. No.: 40068-RP02), mouse monoclonal anti-β-actin antibody (Sigma, cat. No.: A5316), GAPDH (14C10) (Cell Signaling Technology, cat.no.: #2118). Following primary antibody incubation, membranes were incubated with a secondary antibody: biotin-conjugated anti-mouse IgG (Thermo Fisher Scientific, cat.no.: 31800) or anti-rabbit IgG (Thermo Fisher Scientific, cat.no.: A16033). After 1 h incubation at room temperature, the membranes were incubated for 1 h with a tertiary Cy3-conjugated anti-biotin antibody (Jackson Laboratories, cat.no.: 200-162-211). Protein bands were visualized using an Alliance Q9 advanced imaging system (Uvitec).

For the PNGase assay, the PNGase F kit (New England Biolabs, cat. no.: P0704) was used according to the manufacturer’s instructions with a few modifications. The protocol for denaturing conditions was followed and the glycoprotein denaturing buffer was added directly to samples lysed with 4x Laemmli Sample Buffer. The samples were denatured by heating at 100°C for 10 min, they were briefly placed on ice and then centrifuged for 10 sec. GlycoBuffer, 10% NP-40 and H2O were added to the denatured protein lysate according to NEB’s protocol. The reaction mixture was incubated for 1 h at 37°C and then directly subjected to SDS-PAGE and Western blotting.

### Electron microscopy

Cells were fixed with 1.5% (vol/vol) glutaraldehyde in 0.1 M cacodylate buffer (pH 7.4) for 30 min at room temperature, and then stored overnight at 4°C before being exported from the BSL3 facility. Then, the samples were first stained with OsO4 at 4°C for 1 h and subsequently at room temperature with 1% (w/v) uranyl acetate in Milli-Q water for 1 h, with washing steps with 0.1 M cacodylate buffer in between.

After a washing step with Milli-Q water, the samples were dehydrated in increasing concentrations of ethanol (70%, 80%, 90%, 100%), embedded in epoxy resin (LX-112, Ladd Research), and polymerized at 60°C. Sections were collected on mesh-100 copper EM grids covered with a carbon-coated Pioloform layer, and poststained with 7% (w/v) uranyl acetate and Reynold’s lead citrate.

The samples were examined in a Tecnai12 BioTwin or a Twin transmission electron microscope, equipped with an Eagle 4k CCD camera (Thermo Fisher Scientific) or a OneView 4k high-frame rate CMOS camera (Gatan), respectively. For the quantifications, mosaic images of large areas were generated using automatically collected overlapping images (pixel size 2 nm) that were subsequently combined in composite images, as described in (79).. A minimum of 3 large mosaic images from at least 2 different EM grids were analyzed per condition, each one including ∼10-25 cells and DMVs and virus particles were manually annotated and CM regions delineated using Aperio Imagescope software (Leica). On average, each of these large areas contained ∼1,900 DMVs, as well as ∼3,000 virus particles in different stages of assembly. The latter were classified based on their assembly stage: particles in early-stage assembly were defined as those with a budding envelope forming a crescent smaller than a semi-circle, whereas virus particles with a budding envelope forming a crescent larger than a semi-circle or appearing fully budded were assigned to the late-stage assembly class. The perimeter of cells and nuclei in these regions was also manually delineated to calculate densities per µm^2^ of cytoplasm.

### Statistical analysis

Data are represented as mean ± standard deviation (SD) of at least two biological replicates. Statistical significance was calculated using unpaired Student’s t-test, when comparing two groups, or two-way ANOVA and applying multiple comparison corrections, when comparing multiple groups. Statistical significance is denoted with asterisks as: * p<0.05, ** p< 0.01, *** p< 0.001, **** p< 0.0001. Prism version 10 (GraphPad Software Inc.) was used to perform all statistical analyses and calculations.

### Software

Geneious Software (version 10.2) was used for the alignment of partial S and E protein sequences from selected coronaviruses. The list of viruses was selected based on available information from published literature on S and E protein palmitoylation modifications of these viruses. Their protein sequences were then compared to the sequences of MERS-CoV S and E proteins. The following accession numbers were used: MERS-CoV (NC_019843), SARS-CoV (NC_004718) SARS-CoV-2 (NC_045512), MHV (AY700211), TGEV (AJ271965), IBV (NC_001451), HCoV-229E (NC_001451).

Figure annotations (Fig. 5, 7E and S1) and schematics (Fig. 1A) were created using Adobe Illustrator 2025.

## Supplementary data

Available as a separate file.

## Data availability

Raw lipidomics data used to generate and support the conclusions of Figure S2A are available in a public repository (Mendeley Data, doi: 10.17632/sknp5pwj5y.1).

## Acknowledgements

The authors would like to thank Patrick Wanningen, Heidi de Gruyter, and Tim Dalebout for technical assistance, Annelies Boonzaier-van der Laan for confocal microscopy assistance, Nadya Urakova and Quinten Ducarmon for helpful discussions. The study was partially funded by the European Union’s Horizon 2020 research and innovation program under the Marie Skłodowska-Curie grant agreement number 813343 (H2020-MSCA-ITN-813343-INITIATE), and by the Netherlands Organization for Scientific Research (grant NWO-OCENW.M.21.339).

## Author Contribution

Study design, M.S., M.B., and E.J.S.; methodology, M.S., A.W.M.J., and N.B.; investigation, M.S., A.W.M.J., N.B., and A.T.; formal analysis, M.S. and M.B.; resources, M.G., M.B., and E.J.S.; writing – original draft, M.S., M.B., and E.J.S.; writing - editing and final approval, M.S., A.W.M.J., A.T., N.B., M.G., M.B., and E.J.S.; supervision, M.B. and E.J.S.; funding acquisition, M.B. and E.J.S.

