## Supplemental data file for "Perturbation of *de novo* lipogenesis hinders MERS-CoV assembly and release, but not the biogenesis of viral replication organelles"

### Figure S1

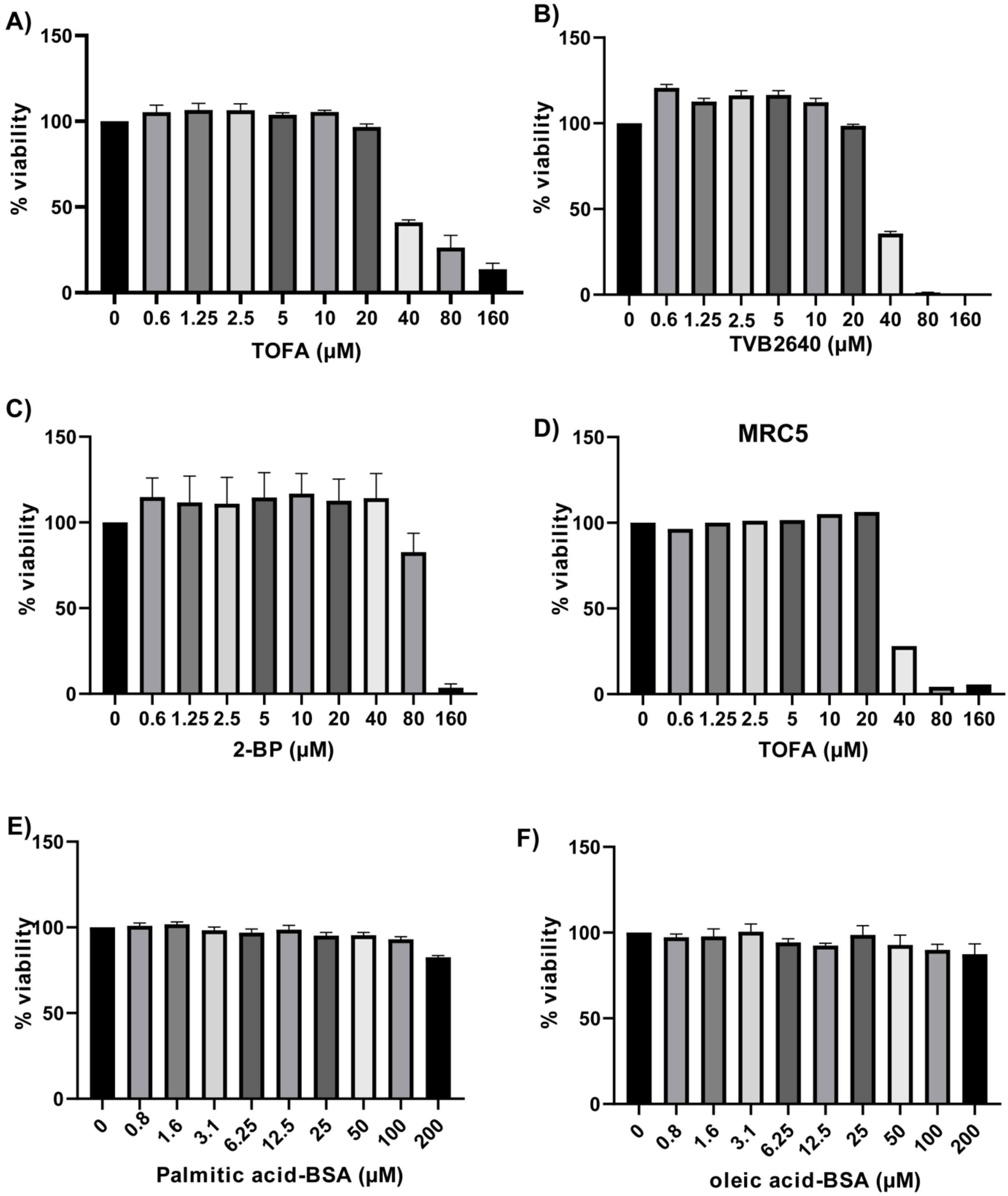


**Figure S1.** Cytotoxicity assays. Cell viability was evaluated following treatment with various small-molecule inhibitors used in this study in different cell lines using an MTS assay at 24 h post compound treatment. Viability was normalized against the DMSO control. A) TOFA in Huh7 cells, B) TVB-2640 in Huh7 cells, C) 2-BP in Huh7 cells, D) TOFA in MRC5 cells, E) palmitic acid-BSA in Huh7 cells, F) oleic acid-BSA in Huh7 cells.

### Figure S2

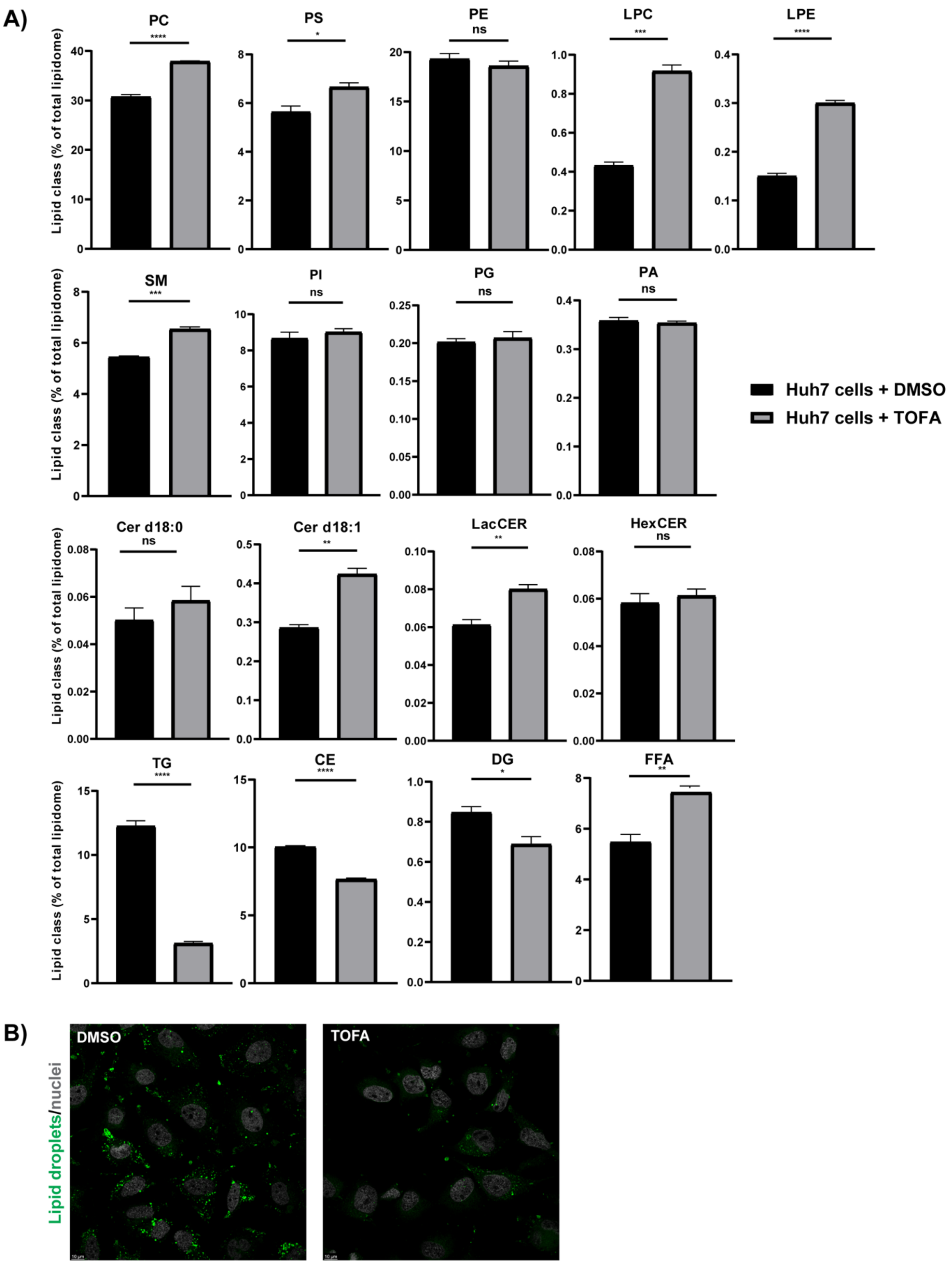


**Figure S2.** Changes in the cellular lipidomic profile upon inhibition of ACC. Huh7 cells were treated with DMSO or TOFA. The amounts of intracellular lipids were determined by mass spectrometry 12 h after compound addition. Different classes of lipids and their relative abundance compared to the total lipidome are depicted in the graphs. A) phosphatidylcholine lipids (PC), phosphatidylserine (PS), phosphatidylethanolamine (PE), Lyso-PC, Lyso-PE, sphingomyelins (SM), phosphatidylinositides (PI), phosphatidylglycerides (PG), phosphatidic acids (PA), ceramides (Cer d18:0), Cer 18:1, lactosyl-CER (LacCER), hexosyl-ceramides, triacylglycerols (TG), cholesterol esters (CE), diacylglycerols (DG) and free fatty acids (FFA). Data are represented as mean ± SD of three biological replicates. Statistical significance was calculated using unpaired Student's t-test, where * $p<0.05$, ** $p< 0.01$, *** $p< 0.001$, **** $p< 0.0001$, B) Huh7 cells were treated as described above and 12h post treatment medium was removed, the cells were washed and incubated with BODIPY 493/503 for 1 h. Cells were then fixed, the nuclei were stained with Hoechst 33342 and the presence of lipid droplets was analyzed by confocal microscopy. Representative microscopy images of two independent biological replicates are depicted.

**Figure S3**

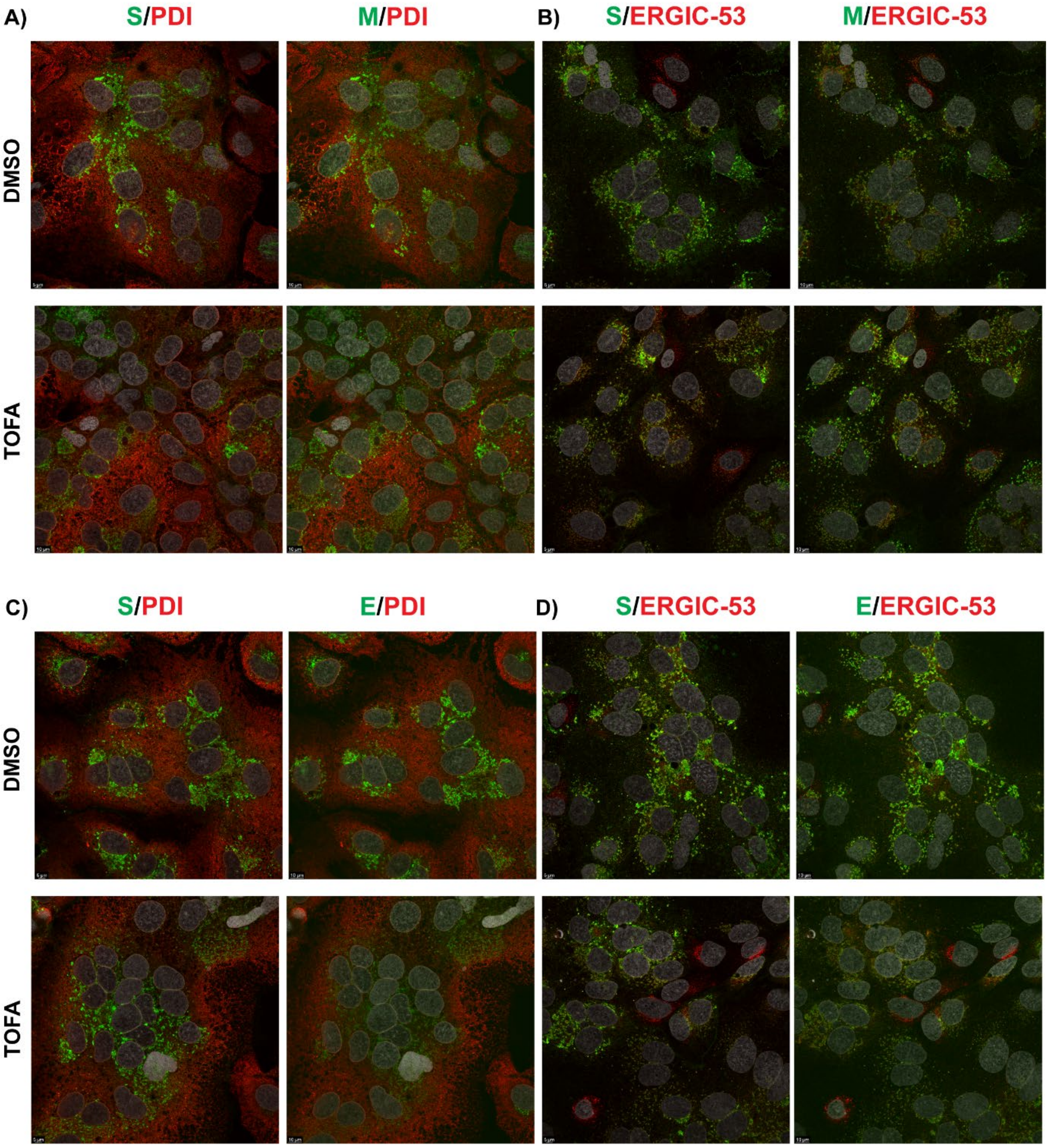


**Figure S3.** Subcellular localization of MERS-CoV structural proteins in the presence of TOFA**.** Huh7 cells were infected with MERS-CoV and treated with TOFA as previously described and fixed at 12 h p.i. Cells were labelled simultaneously for S (green) and a marker for PDI or ERGIC (red) and M or E (green) in combination with a marker for PDI or ERGIC (red). Confocal images are representative of at least two independent biological replicates.

### Figure S4

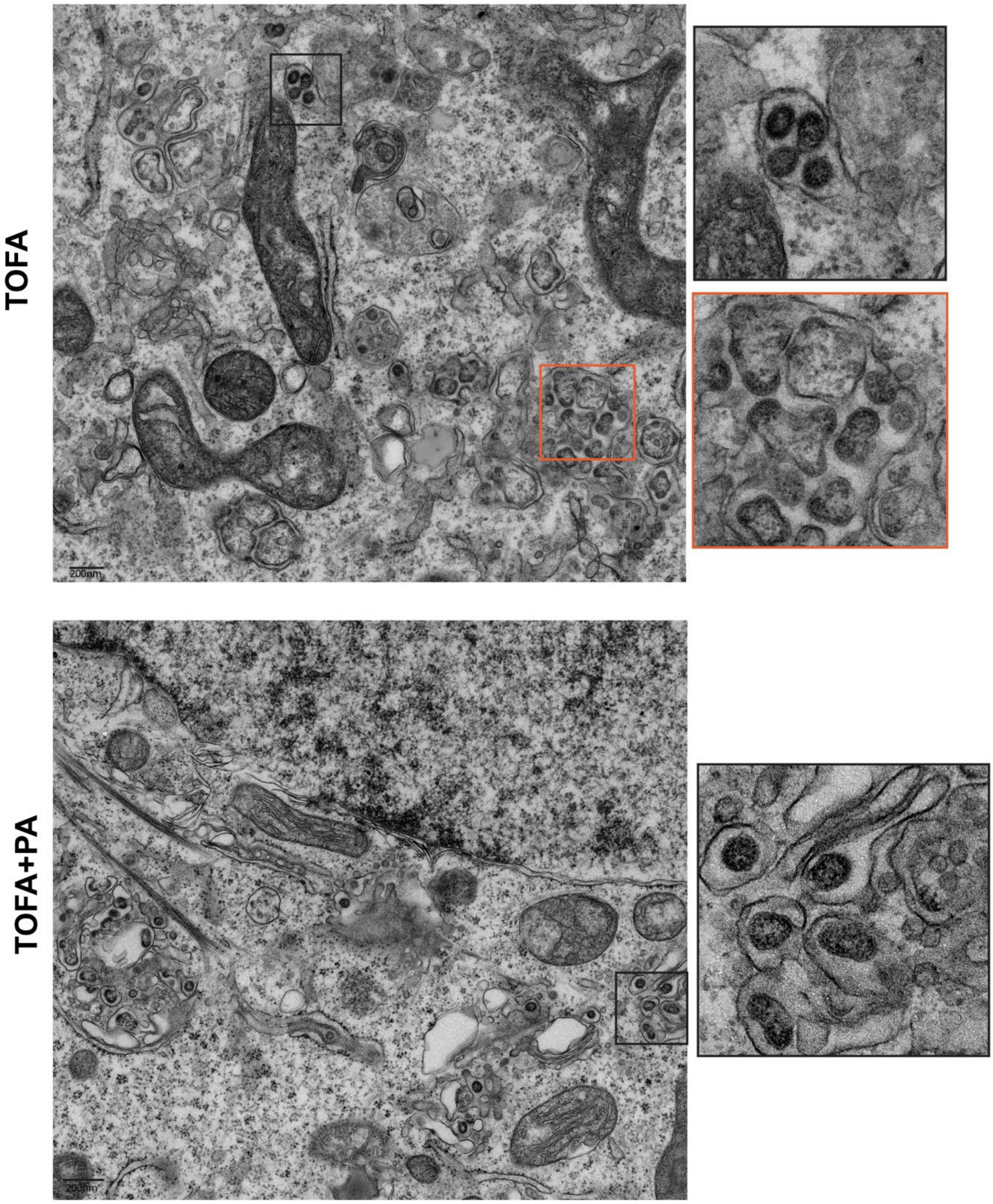


**Figure S4**. Representative EM images of (early and late) assembly events during MERS-CoV replication and TOFA or TOFA and palmitic acid supplementation used for quantifications for Figure 7G. Scale bar is 200nm.

### Figure S5

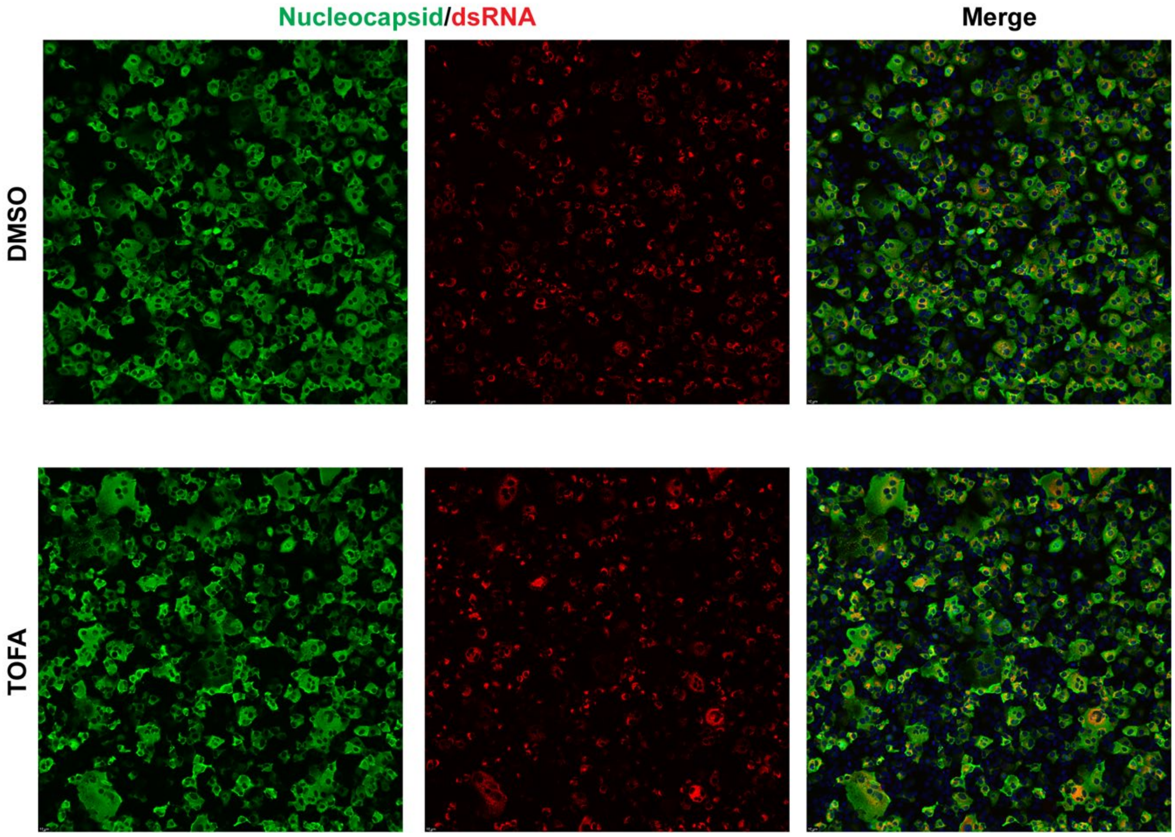


**Figure S5.** Immunofluorescence analysis of MERS-CoV-infected (MOI 5) Huh7 cells, DMSO- or TOFA-treated, 12 h p.i. Cells were labelled for the N protein (green) and dsRNA (red).

**Supplementary Table 1: Primers used in this study**

| Oligonucleotide name | Sequence (5'-3') | Source reference |
|---|---|---|
| ACLY Forward primer | CCTCGAGATCAATCCCCTTGTA | (6) |
| ACLY Reverse primer | CGATGTCACCCCACTTCACTTT | (6) |
| ACC Forward primer | GAGGGCTAGGTCTTTCTGGAAG | (6) |
| ACC Reverse primer | CCACAGTGAAATCTCGTTGAGA | (6) |
| FASN Forward primer | AGCTGCCAGAGTCGGAGAAC | (6) |
| FASN Reverse primer | TGTAGCCCACGAGTGTCTCG | (6) |
| SCD-1 Forward primer | TGCTGCCCACCTCTTCGGATAT | (6) |
| SCD-1 Reverse primer | TAGTTGTGGAAGCCCTCACCCA | (6) |
| DGAT1 Forward primer | GGCATCCTGAACTGGTGTGTG | (6) |
| DGAT1 Reverse primer | GAGCTTGAGGAAGAGGATGGTG | (6) |
| ADRP Forward primer | GGGATCCCTGTCTACCAAGC | (6) |
| ADRP Reverse primer | AGATGTCGCCTGCCATCACC | (6) |
| CPT1a Forward primer | TGAGCGACTGGTGGGAGGAG | (6) |
| CPT1a Reverse primer | GAGCCAGACCTTGAAGTAGCG | (6) |
| ACOX Forward primer | TCCTGCCCACCTTGCTICAC | (6) |
| ACOX Reverse primer | TTGGGGCCGATGTCACCAAC | (6) |
| human b-actin Forward primer | CACAGAGCCTCGCCTTTGC | (73) |
| human b-actin Reverse primer | AATCCTTCTGACCCATGCCC | (73) |
| MERS-CoV N Forward primer | GTACCTCTTAATGCCAATTC | in-house design |
| MERS-CoV N Reverse primer | GAGCCAGTTGCNTTAATTC | in-house design |
| MERS-CoV N probe with TexasRed 5' fluorophore | TCTGTCCTGTCTCCGCCAATAC | in-house design |
| MERS-CoV Nsp2-3 Forward primer | CCGACTCTCTTTAGACTTA | in-house design |
| MERS-CoV Nsp2-3 Reverse primer | ACAGCATGAATGTTGTAC | in-house design |
| MERS-CoV Nsp2-3 probe with FAM 5' fluorophore | AACACTTCTTACAGCAGCAACCTC | in-house design |
